# The transcription factor AP-2β defines active enhancers conferring molecular apocrine cell identity in breast cancer

**DOI:** 10.1101/2025.07.05.663274

**Authors:** Ebtihal Mustafa, Geraldine Laven-Law, Jean Winter, Zoya Kikhtyak, Anthony Bergeron, Gaetan MacGrogan, Amy Dwyer, Stevie M. Pederson, Wayne Tilley, Richard Iggo, Theresa Hickey

## Abstract

The transcription factor Activating Protein 2β (*TFAP2B*; AP-2β) is a candidate marker for the molecular apocrine subtype of breast cancer, which lacks expression of the estrogen receptor alpha (*ESR1*; ER) but sustains a luminal breast cancer phenotype due to the presence of key luminal lineage transcription factors (GATA3, FOXA1) and activity of the androgen receptor (AR). Our objective was to better understand the expression patterns and molecular function of AP-2β in breast cancer. Using multiple clinical cohorts, we show that high *TFAP2B* expression was associated with low proliferation in ER positive (ER+) breast cancers of ductal and lobular histology, but with enrichment for lobular tumours. High *TFAP2B* was also evident in high proliferation tumours lacking ER, with no specific enrichment for lobular histology. Restoring AP-2β expression to highly proliferative ER+ breast cancer cell lines that lack it strongly induced apoptosis. Conversely, reducing AP-2β expression in a molecular apocrine breast cancer cell line model potently inhibited proliferation and cell viability associated with downregulation of MYC oncogene expression. To identify genomic determinants of AP-2β function in molecular apocrine cells in relation to other known transcriptional regulators, we performed chromatin immunoprecipitation followed by DNA sequencing (ChIP-seq) for AP-2β, AR, GATA3, FOXA1, and acetylated lysine 27 in histone 3 (H3K27ac, a marker of active enhancers and promoters). When present alone, AP-2β was preferentially enriched at active promotors. Enhancers bound by AR, GATA3 and FOXA1 were more active when AP-2β was present and genes that define the molecular apocrine phenotype were significantly more likely to have active enhancers co-occupied by all four transcription factors. We conclude that AP-2β plays a context-dependent role in breast cancer and propose that activation of enhancers defining molecular apocrine identity is a key function of AP-2β in the biology of this subtype of breast cancer.

## Introduction

Most human breast cancers fall into one of four well established, clinically relevant molecular subtypes: Luminal A, Luminal B, Her2-enriched (HER2E) and Basal-like (Parker et al. 2009; Perou et al. 2000). All common subtypes of human breast cancer arise from the luminal lineage of mammary epithelial cells, which begins with a luminal progenitor that can differentiate along hormone-sensing or secretory pathways. These subtypes of breast cancer resemble cells at different stages within the luminal lineage (Iggo and MacGrogan 2025; Nguyen et al. 2018; Gray et al. 2022; Reed et al. 2024). Luminal A and Luminal B breast cancers express the estrogen receptor alpha (ER) and resemble epithelial cells within the hormone-sensing lineage, while Basal-like breast cancers lack ER and resemble cells from the secretory lineage (Iggo and MacGrogan 2025). The HER2E subgroup of breast cancers also lacks ER expression but possesses key features of the hormone sensing, not the secretory, lineage. It is now recognized that the HER2E subtype and the luminal androgen receptor (LAR) subtype of triple negative breast cancers (Lehmann et al. 2016; 2011) together form a group we previously named molecular apocrine (MA) breast cancer (Farmer et al. 2005). HER2E, LAR and MA tumours have the same transcriptional profile, defined by expression of FOXA1 and GATA3, key pioneer factors for the hormone-sensing luminal lineage (Jozwik and Carroll 2012), and expression of the androgen receptor (AR), which acts in concert with FOXA1 and GATA3 to transcriptionally regulate luminal differentiation in the absence of ER (Hosseinzadeh et al. 2024; Robinson et al. 2011). We have hypothesized that HER2E/MA/LAR tumours arise from cells within the hormone-sensing lineage that have undergone apocrine metaplasia, a common process as women approach the menopause that reverts breast epithelial cells to their ancestral origin as androgen-driven apocrine (scent) glands (Iggo and MacGrogan 2025; Oftedal 2002; Farmer et al. 2005). Since AR, FOXA1 and GATA3 are expressed in Luminal A, Luminal B and HER2E/MA/LAR tumours, it is likely that other transcription factors are required to determine molecular apocrine identity. We have identified the transcription factor activating protein-2β (*TFAP2B*/AP-2β) as a marker gene for the MA subtype of breast cancer (Farmer et al. 2009; 2005), and hypothesize that it plays a key role in this subtype of disease.

AP-2 transcription factors were first identified in a protein complex binding to the *ERBB2* promoter in Her2-overexpressing mammary cell lines (Bosher, Williams, and Hurst 1995; Hollywood and Hurst 1993). This complex was subsequently shown to contain three related proteins (Bosher et al. 1996), and the family is now known to contain five genes (*TFAP2A-E,* reviewed by (Eckert et al. 2005). AP-2 proteins activate transcription by recruiting CITED family coactivators and the p300/CBP histone acetylases (Braganca et al. 2003; 2002). In ER positive (ER+) breast cancer cells, the AP-2γ isoform regulates *ESR1* gene expression (Woodfield et al. 2007) and cooperates with ER and FOXA1 to regulate the expression of ER target genes (Tan et al. 2011). In normal human breast tissue, AP-2β strongly colocalises with GATA3 and partially colocalises with AR and ER (Raap, Gierendt, Werlein, et al. 2021), pointing to the potential for widespread interactions between AP-2 proteins and these critical hormonal regulators of transcriptional programs in mammary epithelial cells. We have previously reported that *TFAP2B* is a molecular apocrine signature gene (Farmer et al. 2009; 2005), while others have reported that AP-2β is highly expressed by ER+ lobular breast tumours (Raap et al. 2018). Molecular apocrine tumours are high proliferation, poor prognosis tumours that lack ER and are HER2E, whereas the large majority (> 90%) of lobular tumours are low proliferation, good prognosis ER+ tumours that are not HER2E (Arpino et al. 2004). To further understand the potential of *TFAP2B* as a marker for these distinct tumour types, we analysed its mRNA expression across clinical breast cancer cohorts, assessing its expression patterns among different molecular subtypes. We then examined the role of AP-2β as a determinant of cell viability in different breast cancer cell lines and investigated the genomic binding of AP-2β in association with AR, GATA3 and FOXA1 in molecular apocrine breast cancer cells.

## Methods

### Cell culture

The MDA-MB-453, MCF-7, T-47D and 293T/17 cell lines were obtained from the American Type Culture Collection (ATCC, USA). AR knockout T-47D cells were made by CRISPR cas9 cleavage (Chia et al. 2019). All cell lines were cultured at 37°C in a humidified incubator containing 5% CO_2_ in DMEM (MDA-MB-453, MCF-7, 293T/17) or RPMI-1640 (T-47D). Cell lines were regularly screened for the presence of mycoplasma by PCR and authenticated by short tandem repeat profiling (Cell Bank Australia).

### Lentiviral and siRNA transfection

Plasmids and oligonucleotides are listed in Supplementary Table 1. The FUCCI system uses genetically encoded optical probes to monitor cell cycle progression in living cells by fusing fluorescent proteins to regulators of DNA replication origin licensing: Cdt1 is stable in G1, when it loads MCMs onto replication origins; Geminin is stable in S, G2 and M, when it inhibits Cdt1 to prevent re-replication. In the FUCCI CA2 reporter, Cdt1::mCherry stability is regulated by CUL4^DDB1^ and Geminin::Venus stability by APC^CDH1^(Sakaue-Sawano et al. 2017). Lentiviral suspensions were produced as described (Iggo 2022). Cells were transduced at a multiplicity of infection of 1-2 and fully selected with antibiotic before use. Reverse transfection with 2.5-10 nM of siRNA was performed using Lipofectamine RNAiMax (Thermo Scientific) according to the manufacturer’s instructions. Nonspecific siRNA (AllStars Negative Control siRNA, [Qiagen]) was used as a negative control.

### In vitro proliferation assays

The IncuCyte S3 Live-Cell Analysis System (Essen Bioscience, Germany) was used to count live and dead cells after AP-2β overexpression and silencing *in vitro*. Live cell counts were obtained by counting the number of nuclei positive for fluorescent protein expression (mKate2 or neon) conferred by the lentiviral vectors in Supplementary Table 1. Dead cell counts were obtained by counting the number of cells stained with propidium iodide (Sigma), Sytox green (Thermo #S7020) or Caspase-green (Sartorius # 4440). Four images per well were collected with a 10x or 20x objective every 3-6 hours for the periods indicated in the figures and segmented with IncuCyte S3 software.

### Flow cytometry

For cell cycle analysis, cells growing in 6-well plates were trypsinised, fixed in 70% ethanol at 4 °C for >2 hours then incubated with 50 μg/ml propidium iodide, 100 µg/ml RNase A (Sigma), 0.1% Triton X-100 at 37°C for 30 min. Unfixed FUCCI cells were trypsinised and 1 µg/ml DAPI was added to stain dead cells. Data were acquired on FACS Canto II or LSRFortessa X20 flow cytometers (BD Biosciences) at the University of Adelaide Health and Medical Sciences Flow Cytometry Facility and analysed with FlowJo v10.8 (BD Biosciences).

### Western blotting

Western blotting from precast gels (BioRad) was performed as described in reference (Harlow and Lane 1988). Nitrocellulose membranes were probed with the antibodies in Supplementary Table 1 and chemiluminescence was imaged with a Chemidoc (BioRad).

### In situ Proximity Ligation Assay (PLA)

For cell-based assays, MDA-MB-453 cells were seeded on glass coverslips at a density of 3.5 × 10⁵ cells per well in 6-well tissue culture plates containing DMEM supplemented with 5% DCC-stripped FBS. After 48 hours, the cells were treated with either vehicle control (ethanol) or 10 nM dihydrotestosterone (DHT) for 4 hours. The cells were then fixed with 10% neutral buffered formalin for 10 minutes at RT, followed by permeabilization with 0.1% Triton X-100 in PBS for 1 hour. For tumour tissue, sections were cut from formalin-fixed paraffin blocks. Tissue sections and cultured cells were blocked using 10% donkey serum in PBS for 30 minutes at RT in a humidified chamber. Samples were incubated overnight at 4°C with the antibodies listed in Sup Table 1. PLA was performed using the Duolink In Situ Red Starter Kit (Sigma-Aldrich) according to the manufacturer’s instructions. Briefly, species-specific PLA probes were applied for 1 hour at 37°C in a humidified chamber, followed by ligation using a 1:40 dilution of ligase enzyme in ligation buffer for 30 minutes at 37°C. After three washes with 1× wash buffer A, samples were incubated with a polymerase enzyme diluted 1:80 in amplification buffer for 40 minutes at 37°C. Amplified PLA signals were visualized following two 10-minute washes in 1× wash buffer B and a final 1-minute wash in 1:100 diluted wash buffer B. Nuclei were counterstained with DAPI and imaged with an Olympus FV3000 confocal microscope.

**Table 1.** ChIP-seq peaks by location and H3K27 acetylation. The total number of peaks overlapping H3K27 acetylated promoters and enhancers is shown for each transcription factor.

| AR | FOXA1 | GATA3 | TFAP2B | H3K27ac %<br>at<br>All sites | % of<br>acetylated<br>sites at<br>enhancers | Number of<br>Sites<br>(All sites) | Number of<br>sites with<br>H3K27ac | Number of<br>sites at<br>Promoters | Number of<br>sites at<br>Enhancers | Median<br>distance to<br>TSS (bp) |
| --- | --- | --- | --- | --- | --- | --- | --- | --- | --- | --- |
| FALSE | FALSE | FALSE | TRUE | 75 | 11 | 6412 | 4792 | 4273 | 519 | 56 |
| FALSE | FALSE | TRUE | FALSE | 16 | 33 | 4019 | 644 | 433 | 211 | 9268 |
| FALSE | FALSE | TRUE | TRUE | 53 | 28 | 411 | 219 | 158 | 61 | 1807 |
| FALSE | TRUE | FALSE | FALSE | 10 | 38 | 57755 | 5668 | 3541 | 2127 | 12020 |
| FALSE | TRUE | FALSE | TRUE | 54 | 35 | 3967 | 2144 | 1404 | 740 | 2883 |
| FALSE | TRUE | TRUE | FALSE | 12 | 62 | 9906 | 1179 | 443 | 736 | 13653 |
| FALSE | TRUE | TRUE | TRUE | 38 | 58 | 1434 | 551 | 233 | 318 | 8346 |
| TRUE | FALSE | FALSE | FALSE | 10 | 53 | 344 | 36 | 17 | 19 | 10654 |
| TRUE | FALSE | FALSE | TRUE | 49 | 38 | 134 | 66 | 41 | 25 | 2637 |
| TRUE | FALSE | TRUE | FALSE | 13 | 75 | 157 | 20 | 5 | 15 | 13305 |
| TRUE | FALSE | TRUE | TRUE | 48 | 53 | 63 | 30 | 14 | 16 | 4923 |
| TRUE | TRUE | FALSE | FALSE | 12 | 71 | 3580 | 416 | 122 | 294 | 12284 |
| TRUE | TRUE | FALSE | TRUE | 48 | 63 | 1800 | 860 | 322 | 538 | 7779 |
| TRUE | TRUE | TRUE | FALSE | 15 | 81 | 6553 | 1005 | 192 | 813 | 15319 |
| TRUE | TRUE | TRUE | TRUE | 42 | 76 | 6321 | 2674 | 652 | 2022 | 11818 |

All sites means sites where at least one transcription factor is present. Sites designated as enhancers or promoters are by definition acetylated.

### Co-immunoprecipitation

MDA-MB-453 cells were seeded at a density of 6 × 10⁶ cells per 150 mm dish in DMEM supplemented with 10% FBS and cultured for 72 hours. Cells were cross-linked with 1% formaldehyde for 10 minutes, quenched with 0.2 M glycine (pH 7.5), and harvested into PBS containing protease inhibitors. Nuclear extract was sonicated (10 cycles, 30 seconds on/off) using a Diagenode Bioruptor. Lysates were immunoprecipitated overnight at 4°C with 5 µg TFAP2B antibody (Santa Cruz, #SC-390119, Sup Table 1) bound to Protein G Dynabeads (Thermo Scientific, #10004D). The following day, beads were washed and bound proteins were eluted in SDS sample buffer by boiling. Western blot analysis was performed with the antibodies listed in Sup Table 1.

### Chromatin immunoprecipitation sequencing (ChIP-seq)

AP-2β and FOXA1 ChIP-seq experiments were performed as described in reference (Hickey et al. 2021). The AR, GATA3 and H3K27ac ChIP-seq datasets are available in the Gene Expression Omnibus (GSE176010) and described in (Hosseinzadeh et al. 2024). ChIP-seq was performed from three independent replicate experiments with consecutive passages of MDA-MB-453 cells. Cells were seeded at 1 x 10^7^ cells/15 cm plate in normal growth media that was replaced after 24 hours with phenol red-free medium supplemented with 5% Dextran-Coated-Charcoal-stripped FBS. Cells were allowed to grow for 3 days with daily media changes prior to hormone treatment with 10 nM DHT for 4 hours to ensure maximal AR activity in the nucleus. Cells were cross-linked with 1% formaldehyde for 10 min, quenched with 2 mM glycine, washed, pelleted and frozen at -80° C. Cell pellets were lysed and sonicated to produce chromatin fragments of 200-500 bp, and incubated overnight with the primary antibodies in Supplementary Table 1 prebound to protein A or G magnetic beads. Protein-DNA complexes were eluted from the beads and incubated overnight at 65°C to reverse the cross-links. RNA was digested with RNase A for 1 hr at 37°C, protein was digested with Proteinase K for 2 hr at 55°C, and DNA was purified by extraction with phenol-chloroform-isoamyl alcohol followed by ethanol precipitation. Libraries were prepared from 5 ng ChIP-enriched DNA or input DNA using the Qiagen Ultra Low DNA Library Prep Kit and subjected to single-end 75 bp sequencing to a depth ≥20 million reads per sample on an Illumina NextSeq 500 at the South Australian Genomics Centre (Adelaide, Australia).

### Bioinformatics and statistics

Reads were trimmed for adapter sequences and poor-quality bases using the default settings of the Filter by quality tool v1.0.2 on the Galaxy server (Jalili et al. 2020). Raw sequencing files were aligned to GRCh37 using bowtie2 (Langmead et al. 2009). Reads with a minimum mapping quality (MAPQ) <10 and duplicate reads were removed with Samtools v1.8 in Galaxy. ChIP peaks were identified using MACS2 callpeak with default settings (Zhang et al. 2008). The number of reads and peaks in each replicate is given in Supplementary Table 1. Peaks present in at least 2 replicates were retained, yielding 20,317 AP-2β peaks for further analysis after 619 (3.3%) were discarded. Gene annotations were taken from Gencode Release 33. H3K27ac peaks were annotated as promoters if directly overlapping a transcription start site (TSS), or as enhancers otherwise. H3K27ac-HiChIP data was obtained from GSE157381 (Watt et al. 2021) and the vehicle samples were used to call significant interactions using Max-HiC (Alinejad-Rokny et al. 2022). Our published RNA-seq data for MDA-MB-453 cells (GSE176010) (Hosseinzadeh et al. 2024) was used to make present/absent calls for the analysis of the ChIP-seq data. ChIP-seq peaks from AR, FOXA1 and GATA3 and AP-2β ChIP-seq datasets were mapped to genes using H3K27ac-defined regulatory regions and HiChIP interactions as described in the Bioconductor package extraChIPs (https://bioconductor.org/packages/release/bioc/html/extraChIPs.html). Preparatory code for the HiC mapping is available at https://github.com/DRMCRL/MDA-MB-453-H3K27Ac-HiChIP and analytic code for the figures is available at https://drmcrl.github.io/apocrine_signature_mdamb453/. To test for enrichment of apocrine and luminal genes we used the gene lists in reference (Farmer et al. 2009). To find ChIP-seq datasets with similar peak profiles, we analysed consensus AR+GATA3+FOXA1+AP-2β peak sequences on the CistromeDB server (dbtoolkit.cistrome.org). The GIGGLE score is the product of −log10(P value) and log2(odds ratio) from a Fisher test for the number of overlapping peaks in each dataset (Layer et al. 2018; Mei et al. 2017). TCGA and ICGC datasets were downloaded from the Broad and Xena websites, respectively (software.broadinstitute.org/morpheus [BRCA] and xena.ucsc.edu [sp-exp_seq.all_projects.specimen.USonly.xena.tsv]). Normalised gene level expression data for METABRIC were kindly provided by Dr Oscar Rueda (Cancer Research UK, Cambridge University, UK). The EORTC 10994 GSE6861 dataset was downloaded from GEO. The PAM50 classifications were taken from research-pub.gene.com/HER2pancancer/ for TCGA; reference (Rueda et al. 2019) for METABRIC; and calculated with the genefu Bioconductor package for ICGC and EORTC. Normalised RNA-seq data for the pleomorphic lobular dataset was provided by the authors (Bergeron et al. 2021) and classified with the Lobular-Apocrine-Basal LABclassifier R package (https://github.com/ElodieDarbo/LABclassifier) (Bonnefoi et al. 2025). To create the Depmap plot, the 22Q1_Public+Score_Chronos scores for Depmap mammary cell lines were downloaded from the depmap.org website. A one-sided probability was calculated assuming a normal distribution of scores in the breast cancer cell lines. To analyse Incucyte data, cell counts per image were exported as text files and converted to figures in R with ggplot (Wickham 2016). To avoid multiple testing inherent in Incucyte data, Bonferroni-corrected p values were calculated with the lmerTest Bioconductor package (the regression output is given in Supplementary Table 1).

## Results

### *TFAP2B* expression in breast cancer clinical datasets

To explore the paradox that *TFAP2B* has been implicated as a specific marker for both ER-negative molecular apocrine and ER+ lobular breast tumours, we examined *TFAP2B* mRNA expression in multiple clinical datasets. Violin plots of data from the TCGA, METABRIC, ICGC and EORTC breast cancer cohorts grouped by PAM50 subtype show that *TFAP2B* expression is highest in molecular apocrine ("Her2-enriched" in PAM50 terminology) and Luminal A tumours, intermediate in Luminal B tumours, and lowest in Basal-like tumours (Figure 1A; Sup Figure S1A). Scatter plots were used to analyse *TFAP2B* versus *ESR1* expression, with low expression of a proliferation metagene shown in blue, and lobular tumours marked with crosses (Figure 1B). Tumours in the upper right quadrants, which have high expression of *TFAP2B* and *ESR1*, were on average 4.5 times more likely to be lobular than in the other quadrants (n across the four datasets = 3244, of which 10% are lobular), and to have lower expression of the proliferation metagene (more blue symbols). These differences were seen in all four clinical cohorts (Figure 1B; Supplementary Figure S1B) and the reduction in mean proliferation gene expression in the *TFAP2B*-high/*ESR1*-high quadrant compared to the others was highly significant (Supplementary Figure S1C). This suggests that co-expression of *TFAP2B* and *ESR1* marks a subset of ER+ tumours with low proliferative activity and a bias toward lobular histology. In the *TFAP2B*-high/ESR1-low quadrant (upper left), tumours were highly proliferative and not biased toward lobular histology (Figure 1B; Supplementary Figure 1B). Primary ER+ lobular breast cancers can progress to more advanced disease classified as pleomorphic lobular cancer, which can be either ER+ or ER-negative. Because of their rarity, pleomorphic lobular cancers are under-represented in large studies, but analysis of data from a recent study (Bergeron et al. 2021) identified *TFAP2B*-high/*ESR1*-high and *TFAP2B*-high/*ESR1*-low pleomorphic lobular tumour groups of luminal and molecular apocrine molecular subtypes, respectively (Supplementary Figure S1D). Collectively, these data support the concept that *TFAP2B* is a marker of lobular ER+ breast cancer as well as ER-negative molecular apocrine disease of both ductal and lobular histology.

**Figure 1.**
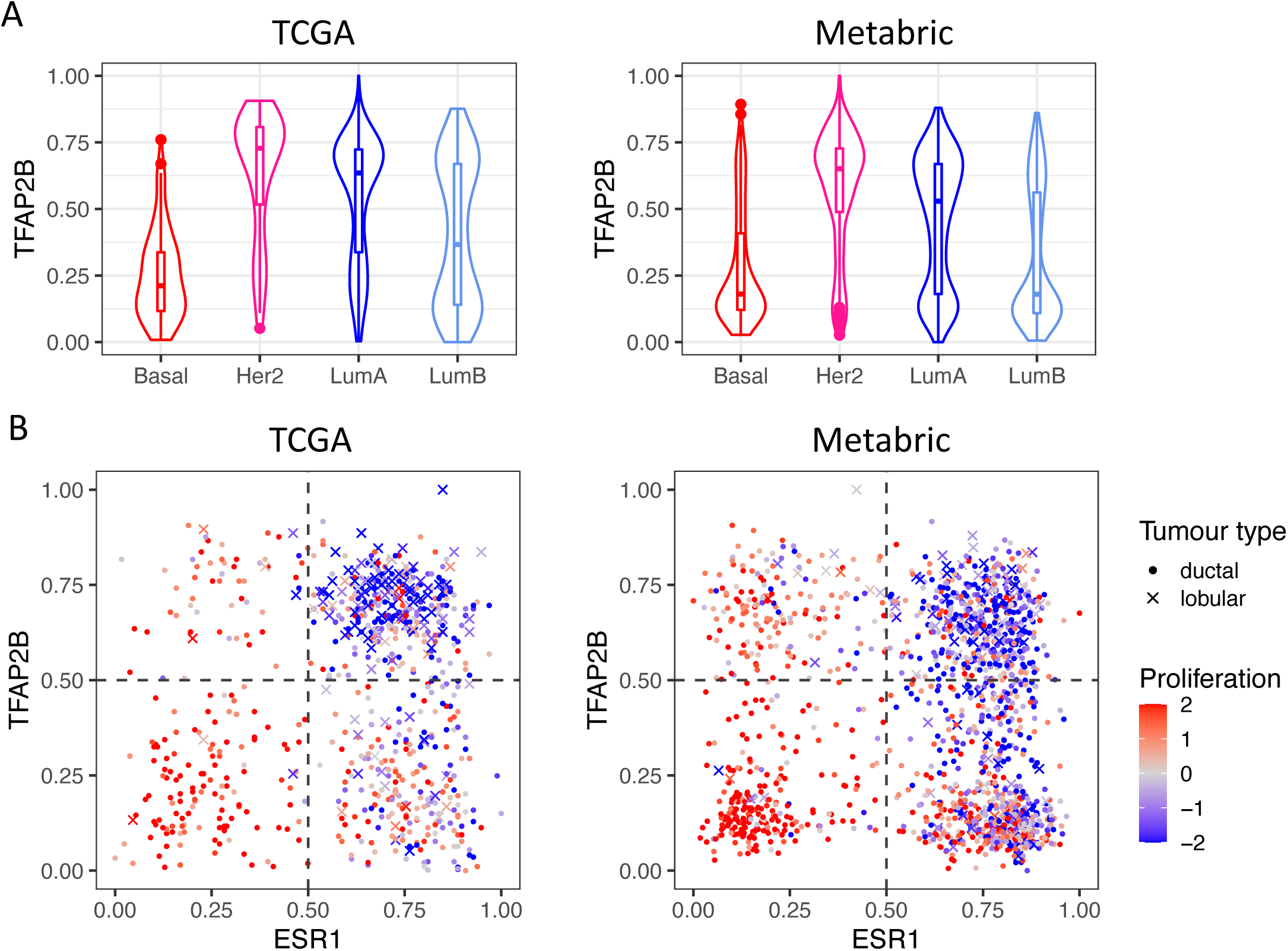
*TFAP2B* expression in the TCGA and METABRIC datasets. A. Violin plots of *TFAB2B* expression in the PAM50 tumour groups. B. Plot of *TFAP2B* vs *ESR1* expression. Lobular tumours are shown with crosses, ductal tumours with dots. A proliferation metagene (the sum of the *TPX2* + *AURKA* + *CCNB2* + *CENPA* gene expression values) was used to colour the symbols, with low proliferation in blue, and high proliferation in red. The expression values are in arbitrary units scaled to the maximum and minimum values in each dataset.

### AP-2β induces apoptosis in ER+ ductal breast cancer cells

Based on the clinical data (Figure 1; Supplementary Figure S1), we predicted that AP-2β expression would not be tolerated in highly proliferative ER+ ductal tumours. To test this, we expressed AP-2β from an inducible vector in two ER+ AP-2β-negative breast cancer cell lines of ductal origin (T-47D and MCF-7, Supplementary Figure S2 A&D). Ectopic AP-2β expression potently reduced proliferation and induced apoptosis in both cell lines (Figure 2A&B, Supplementary Figure S2 B&C), the latter shown by caspase activation and the appearance of a putative caspase-cleaved AP-2β protein fragment (Supplementary Figure S2A). To determine whether death occurred in a particular phase of the cell cycle, we labelled T-47D cells with a FUCCI CA2 reporter (Sakaue-Sawano et al. 2017) that makes live cells red in G1, green in S, and yellow in G2/M. Dead cells were identified with DAPI. Induction of exogenous AP-2β expression led to a G1 cell cycle arrest, with a large proportion of cells dying in G2, as determined by FUCCI labelling and independently validated by flow cytometry (Supplementary Figure S3A&B). Treatment with estrogen did not prevent AP-2β-induced apoptosis (Supplementary Figure S3C). Taken together, these results indicate that DNA synthesis is not required for cell death, and cells die in the G2 phase. We considered several possible explanations for apoptosis induction by AP-2β, such as suppression of ER expression (Supplementary Figure S2A), or absence of driver oncogenes commonly found in ER-negative molecular apocrine cells, like mutant/amplified HER2 and mutant FGFR4. However, ectopic overexpression of ER, mutant ERBB2 or mutant FGFR4 did not alter apoptosis induced by AP-2β in T-47D cells (Supplementary Figure S4A-F). Since molecular apocrine cells have high levels of AR, we tested an *AR* null derivative T-47D cell line and found that AR deletion also had no effect on apoptosis induced by AP-2β (Supplementary Figure S4G&H). We conclude that AP-2β induces apoptosis in ER+ ductal models of breast cancer through a mechanism that is not linked to putative oncogenic mechanisms in molecular apocrine cells.

**Figure 2.**
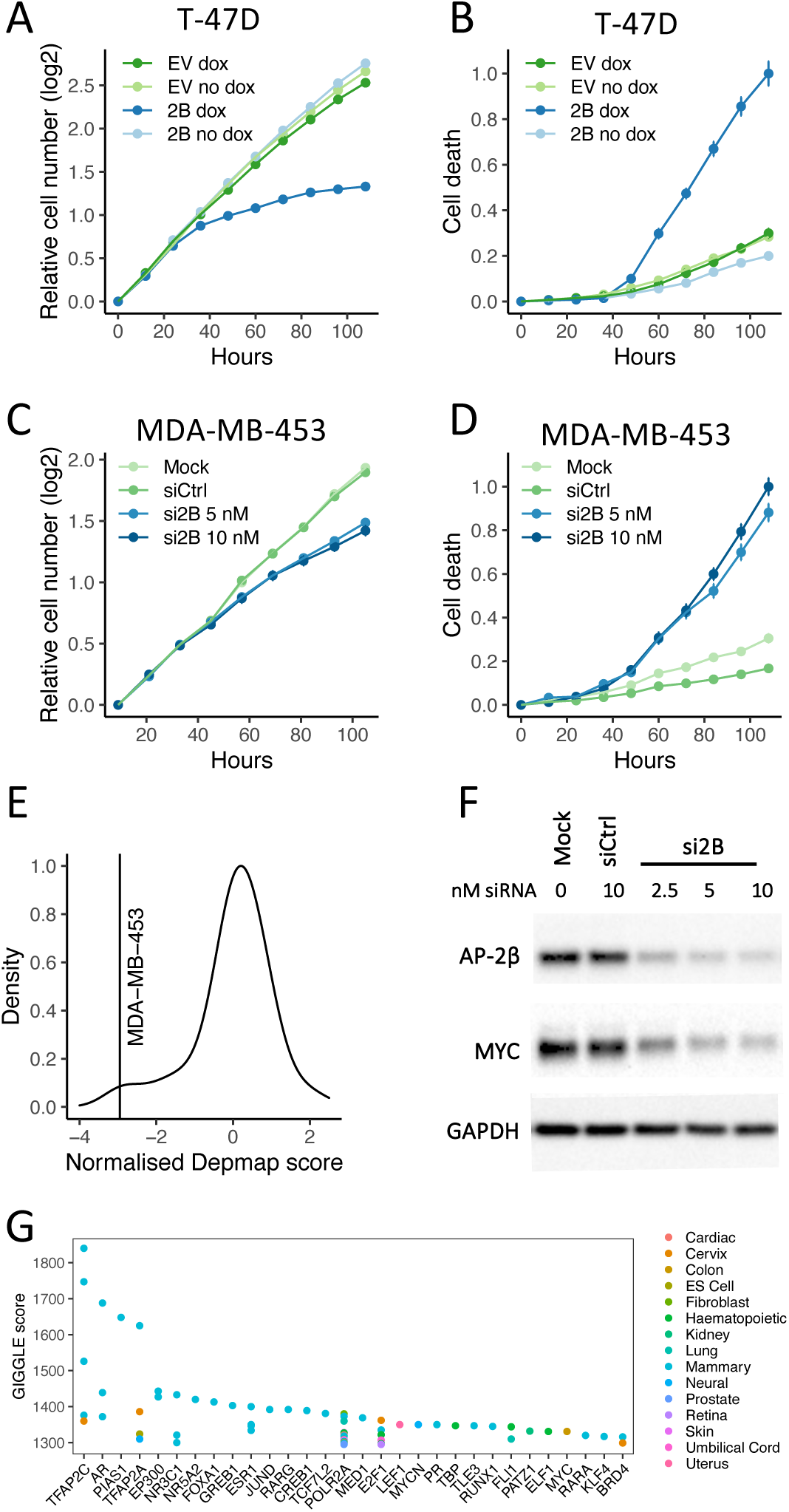
Over-expression and silencing of *TFAP2B* expression in T-47D and MDA-MB-453 cells. A&B. *TFAP2B* expression was induced with doxycycline (dox) in T-47D cells. EV: cells transduced with the empty vector. C&D. *TFAP2B* expression was silenced with siRNA in MDA-MB-453 cells. Cell number was measured by counting nuclei expressing mKate2 protein. Cell death was measured by counting cells stained with sytox-green (B) and caspase-3/7 green dye (D). p values for relevant differences calculated by linear regression are given in Supplementary Table 1 (inhibition of growth and induction of death were highly significant). E. Depmap essentiality scores for *TFAP2B* in breast cancer cell lines. MDA-MB-453 is the most significant outlier (Z = -2.95, p = 0.0016). Note that MDA-MB-453 also expresses the highest level of *TFAP2B* mRNA in the breast cancer cell lines in Depmap. F. Cistrome analysis showing published ChIP-seq studies with peaks overlapping our AP-2β peaks. The GIGGLE score is a composite measure used to rank studies based on the probability of finding overlapping peaks. G. Western blotting showing that the levels of AP-2β and MYC are reduced by siRNA targeting *TFAP2B*.

### AP-2β is required for growth and viability of ER-molecular apocrine breast cancer cells

The most investigated laboratory model of molecular apocrine breast cancer is MDA-MB-453, an ER-negative cell line that expresses high levels of AR and AR target genes (Ni et al. 2011; Doane et al. 2006; Robinson et al. 2011). It is not widely appreciated that MDA-MB-453 cells also represent a model of pleiomorphic lobular breast cancer (Sflomos et al. 2021). This cell line has an *FGFR4* mutation, a feature of metastatic and endocrine-resistant ER+ lobular tumours (Garcia-Recio et al. 2020; Levine et al. 2019), and expresses high levels of AP-2β, making it a perfect model to study tumour cells from the mammary hormone-sensing lineage that have progressed to *TFAP2B+ ESR1-* disease and have a pleomorphic lobular phenotype. *TFAP2B* silencing with siRNA in MDA-MB-453 cells inhibited their growth and induced apoptosis (Figure 2C&D, Supplementary Figure S6), consistent with publicly available CRISPR screen data (depmap.org)(Figure 2E). Since MYC has previously been identified as a potential driver oncogene in MDA-MB-453 cells (Ni et al. 2013), we tested whether silencing of AP-2β expression led to a reduction in MYC expression. This showed that the MYC protein levels fall when AP-2β is silenced (Figure 2F), implicating MYC as a downstream effector. We conclude that *TFAP2B* is critical for the growth MDA-MB-453 molecular apocrine breast cancer cells, implicating it as an oncogene in this context, and that this may occur through induction of MYC expression.

### AP-2β marks active enhancers in ER-molecular apocrine breast cancer cells

To explore potential genomic mechanisms regulated by AP-2β, we performed ChIP-seq to identify AP-2β chromatin-binding events in MDA-MB-453 cells. To identify reproducible binding events, we performed ChIP-seq from three consecutive passages of the cells. The overlap between the peaks identified in each replicate is shown in Supplementary Figure S7. A consensus peak-set (cistrome) was created by centring each peak-set at the peak summit, resizing peaks to 90% of the initial width, then merging overlapping peaks. Comparison of the sequences in our newly generated AP-2β consensus cistrome to ChIP-seq data from other published studies on the cistromeDB server (Layer et al. 2018) showed most similarity with AP-2γ cistromes in breast (mammary) cancer cell lines (MCF-7, MDA-MB-453, BT-474 and SKBR-3) (Figure 2G). Note that the enrichment for breast cancer cell lines in this analysis in part reflects the abundance of ChIP-seq data from these cell lines in the cistromeDB database. The next strongest similarities were with the AR, PIAS1 and AP-2α cistromes in MDA-MB-453 and MCF7 cells (Figure 2G). To assess whether the overlap of the AR and AP-2β cistromes was accompanied by a physical interaction, we examined published mass spectrometry ("AR RIME") data for proteins that interact with AR on chromatin (Hosseinzadeh et al. 2024). This revealed that multiple high-confidence AP-2β peptides were detected in a protein complex with AR on chromatin in MDA-MB-453 cells (Supplementary Figure 7A&B). We confirmed the interaction by co-immunoprecipitation of AR with AP-2β pull-down (Supplementary Figure 7C) and by proximity ligation assay, where the interaction was stimulated by androgen treatment (Supplementary Figure 7D). Proximity ligation assays also detected AR - AP-2β interactions in molecular apocrine tumour tissues (Supplementary Figure 7E), indicating potential clinical relevance.

Next, we examined co-binding of AP-2β and AR with established pioneer factors (FOXA1, GATA3) and acetylated H3K27 (a marker of active enhancers), integrated with chromatin looping data from a published H3K27ac HiChIP study in MDA-MB-453 to link enhancers to promoters (Watt et al. 2021). We first looked for AP-2β peaks at the *MYC* locus, because *MYC* is an AR target gene in this cell line (Ni et al. 2013) and MYC expression was reduced by siRNA targeting *TFAP2B* mRNA (Figure 2F). Analysis of the ChIP-seq data showed that the *MYC* gene is flanked by two active enhancers 65 kb from the gene that are linked to the *MYC* promoter by H3K27ac-marked HiChIP loops, and the flanking enhancers are occupied by AP-2β, AR, GATA3 and FOXA1 (Figure 3A). This indicates that AP-2β, in combination with AR, GATA3 and FOXA1, regulates *MYC* expression in MDA-MB-453 cells, that the reduction in MYC expression after *TFAP2B* siRNA treatment is probably transcriptional in origin, and that the reduced growth and viability of the cells after *TFAP2B* siRNA treatment is likely to be at least partially explained by loss of *MYC* expression. *FGFR4* is a molecular apocrine signature gene (Garcia-Recio et al. 2020; Parker et al. 2009; Farmer et al. 2009) known to be mutationally activated in MDA-MB-453 cells (Roidl et al. 2010). *FGFR4* knockout by CRISPR shows that it is also an essential gene in MDA-MB-453 cells (depmap.org). Our ChIP-seq data show that *FGFR4* is also very likely to be regulated by AP-2β, AR, GATA3 and FOXA1 (Figure 3B), so reduced *FGFR4* expression may also contribute to growth inhibition after treatment of MDA-MB-453 cells with *TFAP2B* siRNA.

**Figure 3.**
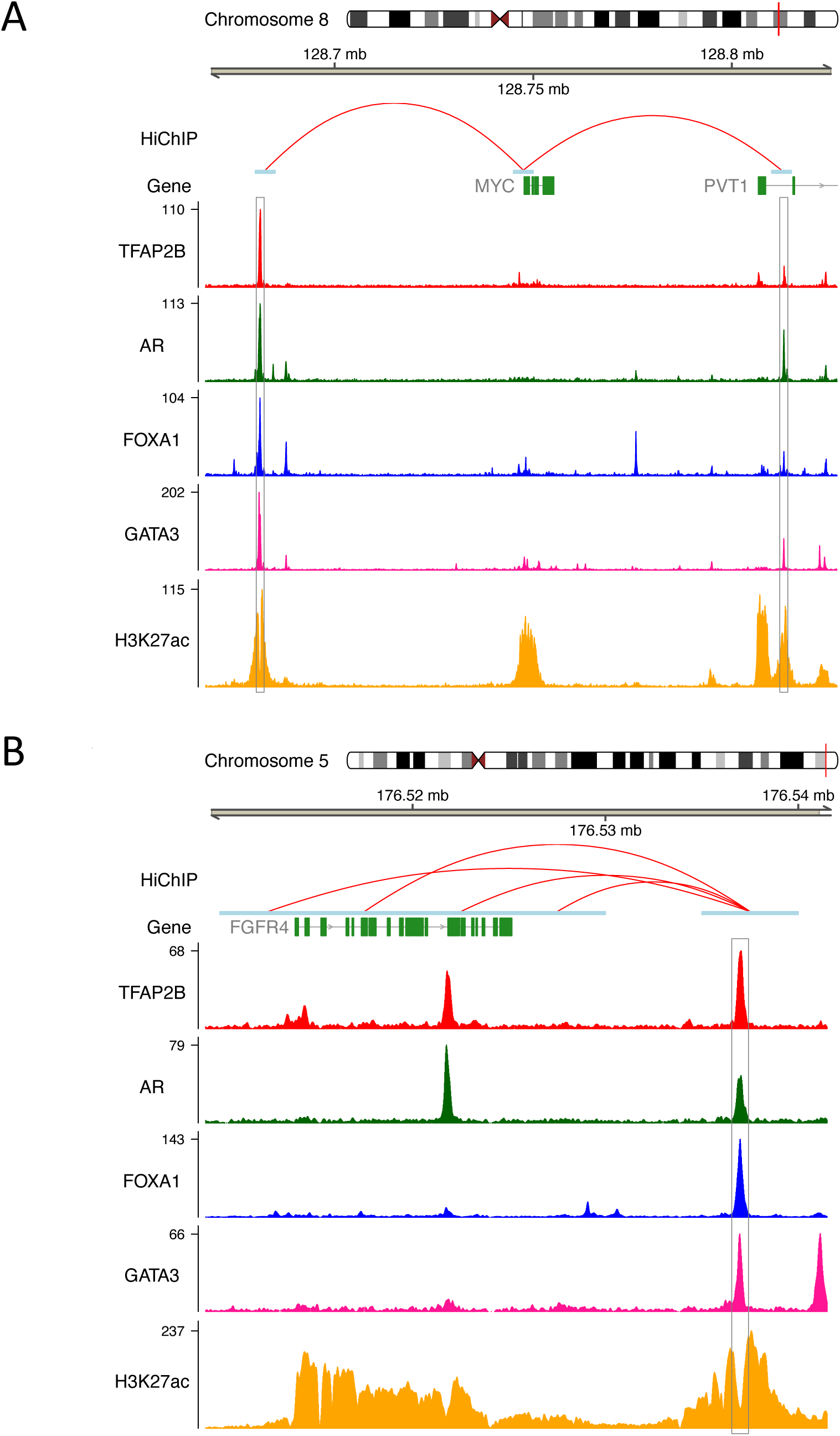
Co-binding of transcription factors and distance from the transcription start site (TSS). A. Upset plot showing the overlap between ChIP-seq peaks for AP-2β, AR, GATA3 and FOXA1. B. Barplots showing the fraction of peaks with respect to the transcription start site (TSS): Core Promoter (<100bp from TSS), Distal Promoter (<2kb from TSS) and Enhancer (>2kb from TSS).

Genome-wide, we observed thousands of enhancers with perfectly aligned peaks for AP-2β, AR, GATA3 and FOXA1 that are flanked by H3K27 acetylation and linked to nearby genes by H3K27ac HiChIP loops (Supplementary Figure S8). They include classic molecular apocrine signature genes (*CLCA2*, *KYNU*, *ERBB2*; Supplementary Figure S8A-C), and the genes encoding the ChIP-seq factors themselves (*TFAP2B*, *FOXA1*, *AR, GATA3*; Supplementary Figure S8D-G). Binding by all four transcription factors (AR, FOXA1, GATA3, AP-2β) occurred at 6321 sites, far above the background expectation (<1 site by chance if the factors bound to DNA independently of one another), indicating coordinated transcriptional regulation (Figure 4A and Table 1). Of these co-bound sites, 2022 mapped to enhancers and 652 to promoters (Table 1). Enhancer targeting increased with the number of co-bound transcription factors (Figure 3B) and AP-2β substantially increased the total number of enhancer sites with multi-transcription factor occupancy (2011 with vs. 813 without AP-2β). Overall, the fraction of such sites classified as enhancers (Figure 3B) was slightly lower in the presence of AP-2β presumably because AP-2β collaborates with other factors at promoters (Figure 3B, Table 1). These findings suggest that AP-2β not only increases enhancer occupancy when acting in concert with AR, GATA3 and FOXA1, but also acts independently of these factors at promoters, probably in cooperation with transcription factors not examined here, such as JUND, RARG, CREB1, TCF7L2, LEF1 and E2F1, whose cistromes overlap with that of AP-2β (Figure 2F).

**Figure 4.**
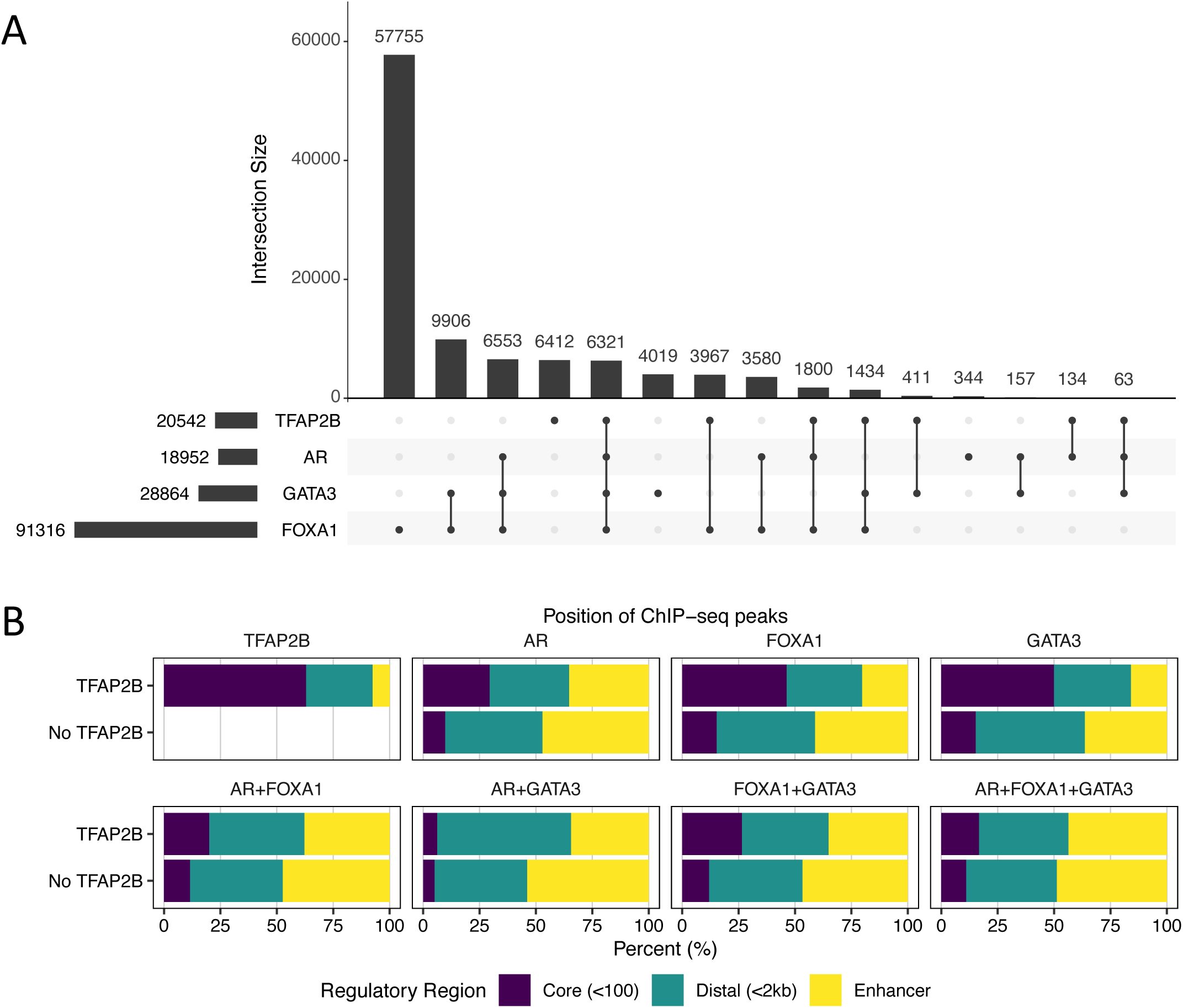
ChIP-seq peaks at enhancers linked to the *MYC* and *FGFR4* genes. ChIP-seq for AP-2β, AR, FOXA1, GATA3 and H3K27 acetylation at enhancers near the *MYC* (A) and *FGFR4* (B) genes. The red arcs show H3K27 acetylation HiChIP loops connecting enhancers to their target genes.

To distinguish between active and inactive genomic regions, we looked for H3K27ac peaks flanking the transcription factor peaks. The probability of a transcription factor peak being flanked by an H3K27ac peak was always higher when AP-2β was present (Figure 5A). When those active sites were split into promoters and enhancers, AP-2β alone (ie, in the absence of AR, GATA3 or FOXA1) showed a strong preference for binding to promoters (Figure 5A). Strikingly, 75% of peaks with AP-2β alone were flanked by acetylated H3K27, vastly more than was seen with any other single factor (10-16%, Table 1). Increased H3K27 acetylation in the presence of AP-2β is clearly visible in heatmaps of aligned peaks (Figure 5B&C lower panels), and in the corresponding H3K27 acetylation density plots (Figure 5B&C top right panels). In Figure 5B the peaks were selected at random (Control) or at sites bound to a single factor (AR, FOXA1, GATA3 or AP-2β). Note that since we can only select among peaks called by MACS2, the Control is not enriched for any particular factor but it is not random DNA, so we do expect to see some peaks in the Control heatmaps. In Figure 5C, instead of asking for binding by single factors, we asked whether AP-2β increases acetylation at sites bound to AR, GATA3 and FOXA1: the lower panels are for unselected sites, the upper panels show enhancers. Since the enhancers were defined by H3K27 acetylation, we expect a higher background than at randomly chosen sites. In all cases, the density plots in the top right panels show that the presence of AP-2β was associated with an increase in the degree of H3K27 acetylation. We conclude that AP-2β is an important determinant of H3K27 acetylation at enhancers and promoters in MDA-MB-453 cells. This is shown schematically in the model depicted in Figure 5D: when AR+FOXA1+GATA3+AP-2β bind to enhancers, the flanking nucleosomes are likely to be acetylated on H3K27. The simplest explanation is that p300/CBP recruited by AP-2β creates a "poised" state allowing for rapid activation of p300 recruited by AR in response to hormonal signals.

**Figure 5.**
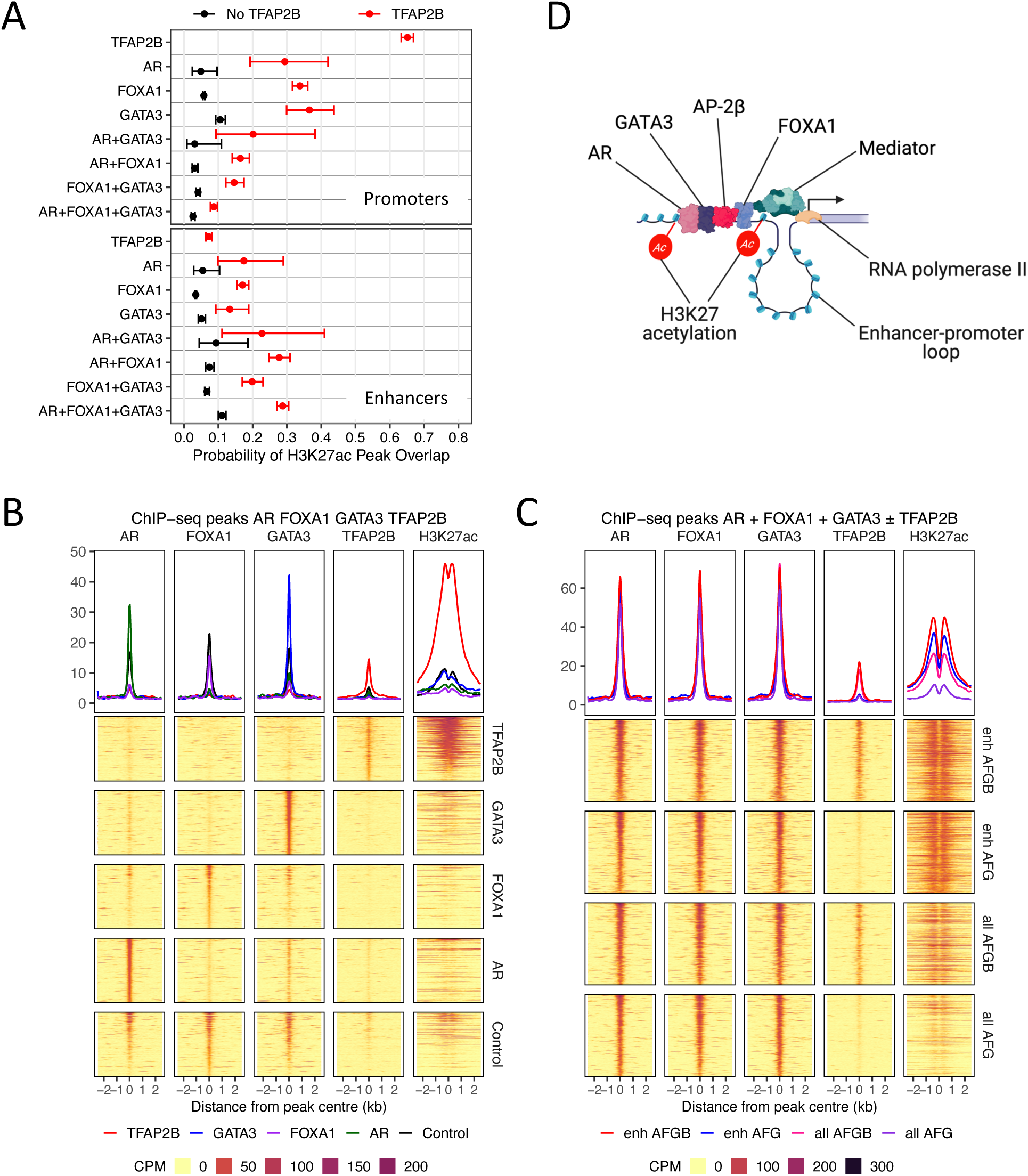
AP-2β promotes H3K27 acetylation. A. The probability of H3K27 acetylation at ChIP-seq peaks bound by AR, GATA3 and FOXA1 alone or together is higher when AP-2β is also present. B&C. Heatmaps showing aligned peaks. In B, the peaks were selected at random (Control), or for individual factors. C shows peaks bound to AR+FOXA1+GATA3 ("AFG") or AR+FOXA1+AP-2β ("AFGB") either at unselected sites ("all") or at enhancers ("enh"). The top right density plots show that there is always more H3K27 acetylation when AP-2β is present. D. Model showing H3K27 acetylation at enhancers when AP-2β is present.

### AP-2β, AR, FOXA1, GATA3 and H3K27 acetylation predict mammary cell type

To test whether the co-binding of AR, FOXA1, GATA3 and AP-2β in MDA-MB-453 cells defines molecular apocrine cell identity, we investigated whether sites bound by all three factors are more likely to be present near molecular apocrine signature genes. We previously defined these genes by "in silico microdissection" (Farmer et al. 2009): we used multiple regression to explain the expression of each gene in a breast cancer dataset as a linear combination of "prototypic" genes that are markers for common cell types and processes in breast cancer. The coefficients in the linear model were then ranked to make signatures for each cell type or process (Farmer et al. 2009). The prototypes were chosen to represent the major clusters in breast microarray data: proliferation, hypoxia, interferon and cell type (luminal, molecular apocrine, stroma, fat, T cell, macrophage) (Farmer et al. 2009; Perou et al. 2000). To test whether transcription factor binding and/or chromatin modification can predict molecular apocrine cell identity we therefore asked whether the signature genes in the linear model have ChIP-seq peaks for AR, FOXA1, GATA3 and AP-2β, with or without flanking H3K27ac peaks. Table 2 shows that all four transcription factors plus H3K27ac gave a highly significant enrichment for apocrine signature genes. This was true at both promoters and enhancers (Table 2). Inclusion of GATA3 gave a weak prediction for luminal cells, consistent with its higher expression in ER+ tumours. We conclude that H3K27-acetylated enhancers and promoters occupied by AR, FOXA1, GATA3 and AP-2β play an important part in defining molecular apocrine cell identity.

**Table 2.**
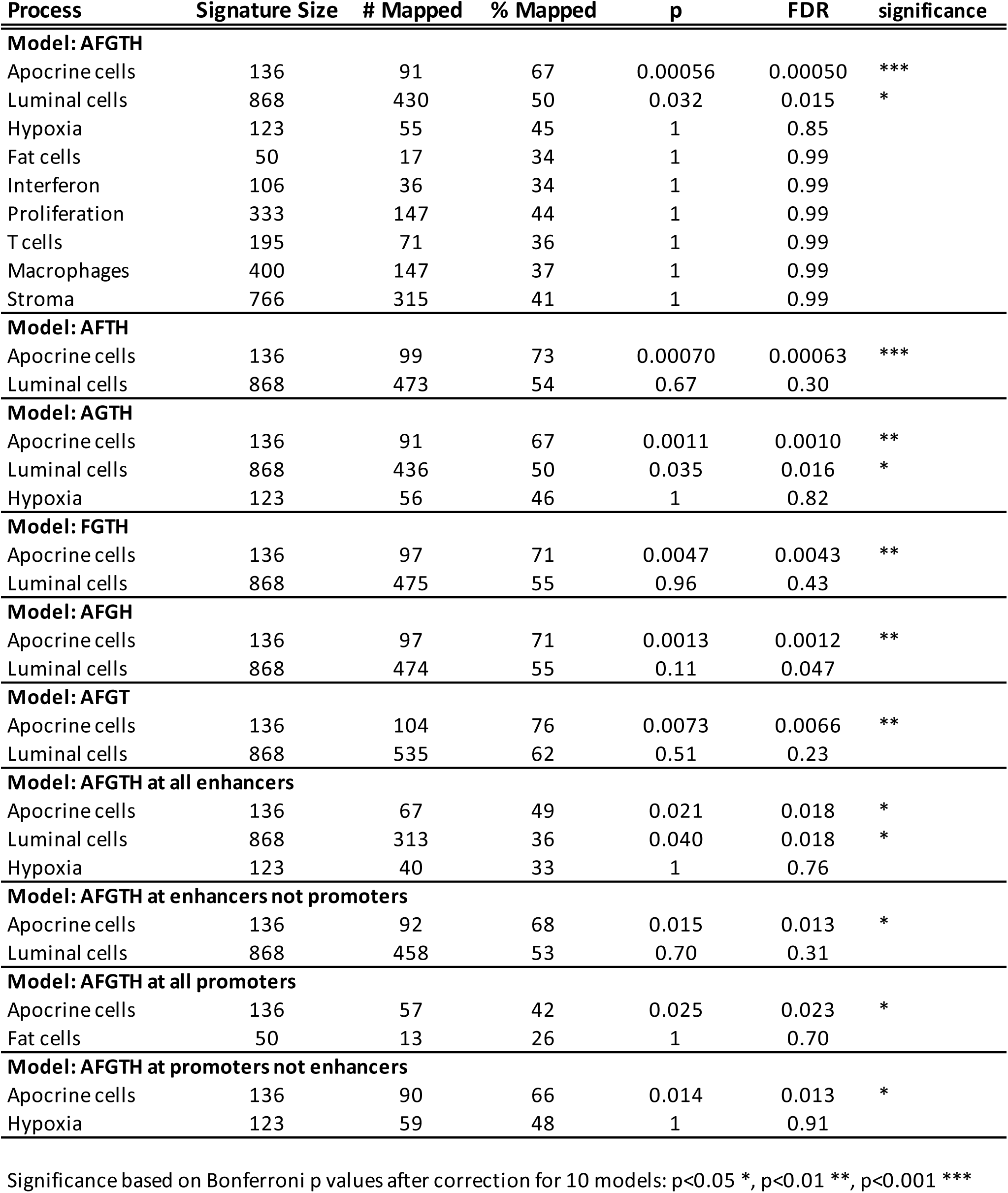
Enrichment for genes expressed by particular cell types or in particular states. *In silico* microdissection of breast cancer microarray data was performed by multiple regression (Farmer et al. 2009) to create signatures for each cell type or process. The number of genes in each signature overlapping peaks for the indicated factors was compared to randomly selected genes by Fisher test. Significant results are shown in bold font. Results for all of the processes (cell type, etc) are shown for the full model; only the best non-significant result is shown for the models with fewer factors. The factors included in the model are abbreviated as A (AR), F (FOXA1), G (GATA3), T (AP-2β), H (H3K27 acetylation). Some genes have peaks at both enhancers and promoters; models with "all enhancers" and "all promoters" incudes these genes; models with "promoters not enhancers" and "enhancers not promoters" exclude these genes.

### Motif and cistrome analysis of molecular apocrine enhancers

Transcription factors can be recruited to DNA either by direct contact or through bridging proteins. To determine whether DNA binding was direct, we examined the sequences within the ChIP-seq peaks for the presence of known motifs for AR, FOXA1, GATA3 and AP-2β (Supplementary Figure S9A). Unsurprisingly, these motifs were strongly enriched in their cognate ChIP-seq peaks, indicating that DNA binding is likely to be direct (Supplementary Figure S9B). To complement the motif analysis, we performed cistrome analysis comparing our peaks with ChIP-seq peaks in published studies. We screened for cistromes matching our AR+FOXA1+GATA3+AP-2β cistrome with and without flanking H3K27ac peaks (Supplementary Figure S10A&B) and our AR+FOXA1+GATA3+AP-2β cistrome in enhancers and promoters (Supplementary Figure S10C&D). Compared to cistrome analysis with AP-2β alone, which produced hits to a wide range of cell types (Figure 2F), all of the top hits with AR+FOXA1+GATA3+AP-2β were to mammary and prostate cell lines and tissues, with a large preponderance of mammary cells (Supplementary Figure S10). The hits were from ChIP-seq studies with classic transcriptional regulators in mammary and prostate cells: nuclear receptors (NR3C1/Glucocorticoid Receptor, PR, AR, ER), their binding partners (FOXA1, GATA3, TCF7L2, AP-2α, AP-2γ), coactivators (p300, PIAS1, GREB1) and corepressors (TLE3, SIN3A, HDAC2). Comparing active with inactive sites, GREB1 (Mohammed et al. 2013) was only present in the active list, and SIN3A and HDAC2 were only present in the inactive list, consistent with their roles as coactivators and corepressors, respectively. TCF7L2 recruits TLE3 to repress transcription at FOXA1 sites (Ni et al. 2013). TCF7L2 was only present at inactive sites, and TLE3 had a higher GIGGLE score at inactive sites, supporting a role for a FOXA1-TCF7L2-TLE3 complex at inactive sites. p300 was only seen at active sites, consistent with its role as an H3K27 acetyltransferase. Collectively, these findings indicate AP-2β acts in concert with the AR, GATA3 and FOXA1 at enhancers in molecular apocrine cells, and acts mainly in concert with other transcription factors at promoters.

## Discussion

In this study, we have functionally implicated AP-2β as a critical driver of molecular apocrine breast cancer. We have shown that AP-2β expression in clinical cohorts is highest in luminal A and molecular apocrine tumours, including lobular tumours. AP-2β expression is low in most luminal B tumours, and we show that reintroduction of AP-2β into ER+ cell lines slows their growth and induces apoptosis. Molecular apocrine tumours have the opposite phenotype, namely loss of ER expression and retention of AP-2β expression, and we show by silencing its expression that AP-2β is required for normal growth of these cells. The most important conclusion from this study is that enhancers bound by AP-2β, AR, GATA3 and FOXA1 are significantly more active (i.e., flanked by acetylated H3K27) at genes that define molecular apocrine identity and represent a core transcriptional complex important for this lineage. We have shown that AP2-β binds to mammary enhancers alongside two pioneer factors critical for the luminal lineage, FOXA1 and GATA3, in addition to a hormone signal transducer, AR. The probability of H3K27 acetylation was much higher in the presence of AP-2β. While the other factors also increase H3K27 acetylation, the effect of AP-2β was much greater. AP-2 proteins activate transcription by recruiting CITED family coactivators and p300/CBP (Braganca et al. 2002; 2003). p300 and CBP have multiple transcription factor interaction sites, but their HAT activity is regulated by an autoinhibitory peptide that blocks the active site. Cross-acetylation by another p300/CBP molecule is required to lift the autoinhibitory loop out of the active site (Delvecchio et al. 2013). A possible model to explain our results is that AP-2β maintains enhancers in a poised state. When AR binds to those enhancers in response to androgen, the resident p300/CBP recruited by AP-2β could cross-acetylate the autoinhibitory loop in the incoming p300/CBP recruited by AR, leading to rapid activation of transcription.

When AP-2β binds to DNA in the absence of AR, GATA3 and FOXA1, it binds preferentially to promoters that are strongly marked by H3K27 acetylation. Cistrome analysis (Figure 2F) shows that it probably cooperates with transcription factors like CREB1, TCF7L2, E2F1, RUNX1, FLI1, ELF1 and MYC at these sites, and it does so in a wide variety of different tissues. We have focused our attention on determinants of mammary hormone sensing cell identity, but there were more promoters to which AP-2β binds in the absence of AR, GATA3 and FOXA1 than there were enhancers to which AP-2β binds alongside AR, GATA3 and FOXA1 (Table 1). Some of the pathological phenotypes of perturbed AP-2β expression could result from interference with its function at the promoter. Indeed, an effect at promoters might better explain the diverse and contradictory roles which have been assigned to AP-2β in different tumour types (Raap, Gierendt, Kreipe, et al. 2021). Our results leave open the possibility that drugs that induce AP-2β expression could be used to attack ER+ tumours that do not express AP-2β, or that drugs that inhibit AP-2β could be used to attack ER-tumours that do express AP-2β, but phenotypic plasticity might quickly subvert the response.

In the context of ER-negative breast cancers, *TFAP2B* is almost exclusively expressed by molecular apocrine tumours, with little or no expression in basal-like tumours (Farmer et al. 2009), but it is also an established marker for ER+ lobular tumours (Raap et al. 2018). In agreement with Raap and colleagues (Raap, Gierendt, Werlein, et al. 2021; Raap, Gierendt, Kreipe, et al. 2021), our results point to AP-2β being a marker for mature luminal differentiation in ER+ tumours. We suggest that AP-2β is expressed by well differentiated luminal cells but lost as ER+ cells become more transformed. The physiological role of normal ER+ mammary hormone-sensing cells is to convert systemic endocrine signals into paracrine growth factor signals that stimulate the proliferation of neighbouring ER-negative cells (Beleut et al. 2010; Clarke et al. 1997). This explains the pattern of gene expression shown in Figure 1: low proliferation tumours tolerate the presence of both ER and AP-2β, but high proliferation tumours express only one of them, giving rise to either ER+ AP-2β- luminal B tumours, or ER- AP-2β+ molecular apocrine tumours. In the former, we have shown that forced expression of *TFAP2B* induces apoptosis. We note that this is not true of all ER+ cell lines, some of which retain AP-2β expression, albeit at very low levels (for example, ZR75-1). Conversely, in molecular apocrine tumours, *TFAP2B* may acquire the oncogenic functions that have been ascribed to it in other malignancies, notably alveolar rhabdomyosarcoma (aRMS), a form of sarcoma caused by *PAX3-FOXO1* translocations (Ebauer et al. 2007). In aRMS, the PAX3-FOXO1 transcription factor transactivates *FGFR4* and *TFAP2B* transcription. *FGFR4* is a molecular apocrine signature gene (Garcia-Recio et al. 2020; Parker et al. 2009; Farmer et al. 2009), whose kinase domain is activated by mutations in lobular tumours (Levine et al. 2019), as it is in aRMS (Taylor et al. 2009). The MDA-MB-453 breast cancer cell line appears to phenocopy aRMS: it has high level expression of AP-2β and an activating mutation in *FGFR4* (Bajrami et al. 2018; Garcia-Recio et al. 2020).

Consistent with the behaviour of *TFAP2B* in aRMS, silencing of *TFAP2B* expression in MDA-MB-453 cells reduces *MYC* expression and induces apoptosis. MDA-MB-453 is a molecular apocrine cell line with pleomorphic lobular histology and high expression of wild type *ERBB2*. Pleomorphic lobular tumours commonly have *ERBB2* amplification or *ERBB2* activating mutations (Riedlinger et al. 2021), and 60% have apocrine features (Bergeron et al. 2021). Unlike high grade ductal tumours, high grade lobular tumours can tolerate the presence of both ER and AP-2β (Bergeron et al. 2021). This could be related to the presence of molecular apocrine oncogenes in the cells, but activated mutants of *FGFR4* and *ERBB2* did not protect T-47D cells from death when we overexpressed *TFAP2B* in them.

In conclusion, the AP-2β transcription factor binds to the enhancers and promoters that define molecular apocrine identity. It cooperates with AR, GATA3 and FOXA1 at these sites, where it promotes H3K27 acetylation. We propose that a key role of AP-2β in breast cancer is activation of enhancers that define the mammary sensory cell lineage.

## Accession numbers

Chip-seq and RNA-seq GSE176010

## Acknowledgements

We thank Prof. Jason Carroll for commenting on an early version of the manuscript.

## Author Contributions

EM, GLL, ZK, and RI performed experiments; GMG and AB provided clinical data; EM, GLL, RI, AD, SP, WT, TH analysed or interpreted data; TH, JW, AD, WT and RI designed the study and supervised the work; EM, RI and TH wrote the manuscript. All authors commented on the manuscript and approved the final version.

## Figures and Tables

**S1. *TFAP2B* expression in the ICGC, EORTC 10994 and Dijon datasets.**

A. Violin plots of *TFAB2B* expression in the PAM50 tumour groups.

B. Plot of *TFAP2B* vs *ESR1* expression. Tumours with low *CDH1* expression (lobular tumours) are shown with crosses, other tumours with dots. A proliferation metagene was used to colour the symbols, with low proliferation in blue, and high proliferation in red. The expression values are in arbitrary units scaled to the maximum and minimum values in each dataset.

C. Proliferation in the quadrants shown in Figure 1B. The mean expression and SEM are plotted for a proliferation metagene in tumours with high *ESR1* and *TFAP2B* expression (UR, upper right quadrant); high *ESR1* and low *TFAP2B* (LR quadrant); low *ESR1* and high *TFAP2B* (UL quadrant); low *ESR1* and low *TFAP2B* (LL quadrant). A. TCGA; B, ICGC; C, Metabric; D, EORTC datasets. Pairwise comparison of UR to the other quadrants gives p<0.00001 for all datasets (ANOVA/Tukey).

D. *TFAP2B* vs *ESR1* expression in high grade lobular and ductal tumours. Low proliferation/grade 1 tumours were not included in this study. The dots were coloured by LAB class (Bonnefoi et al. 2025). The basal-like/ductal tumours were included in the Dijon study as a control group (they are not lobular). The differences in the fraction of lobular tumours between datasets reflect the priorities of the clinical groups that performed the studies, not the population frequency of the tumours.

**S2. AP-2β expression in T-47D and MCF7 cells.** A. Western blot for AP-2β and ER in T-47D cells after induction of AP-2β expression with doxycycline. The lower AP-2β band in the final lane is probably cleaved at amino acid 28 in the sequence "DRHD". The equivalent site in AP-2α is an experimentally proven caspase cleavage site (Nyormoi et al. 2001). B. Incucyte growth and C. death curves in MCF-7 cells after induction of AP-2β expression with doxycycline. Cell number in panel B was measured by counting nuclei expressing neon protein. Cell death in panel C was measured by counting cells stained with propidium iodide. p values for relevant differences calculated by linear regression are given in Supplementary Table 1 (inhibition of growth and induction of death were highly significant). D. Western blot for AP-2β and ER in MCF-7 cells after induction of AP-2β expression with doxycycline.

**S3. Cell cycle analysis showing apoptosis induction by AP-2β overexpression in T-47D cells.** A. FUCCI analysis. The colour of dead cells was used to infer the cell cycle phase in which they died. The cells contain an integrated FUCCI CA2 reporter. DAPI was used to identify dead cells, with live cells in the plots coloured blue and dead cells coloured red. AP-2β expression was induced with doxycycline. B&C, Barplots showing cell cycle distribution and apoptotic fraction. Cells were grown in full medium (FBS) in panels B&C, and in stripped serum minus (NS) or plus 1 nM estradiol (E2) in panel C. Induction of *TFAP2B* expression with doxycycline led to cell death, regardless of the basal proliferation rate.

**S4. Apoptosis induction by AP-2**β **in T-47D cells is not suppressed by exogenous *ER*, *ERBB2*, *FGFR4* or deletion of *AR*.** A-F. *ER*, *ERBB2* and *FGFR4* were expressed from a constitutive lentiviral vector and *TFAP2B* expression was induced with doxycycline: A&B, *ESR1;* C&D, *ERBB2* containing the activating mutation D769Y; E&F. *FGFR4* containing the activating mutation Y376C. G&H. *TFAP2B* expression was induced with doxycycline in *AR*-knockout cells. A, C, E, G. Incucyte curves showing dead cells stained with propidium iodide, sytox green or caspase-green, as indicated. Cell death was normalised to the maximum value after *TFAP2B* expression and represents the number of dead cells in A-D, and the ratio of dead cells to live cells in E. B, D, F. Western blots showing expression of the transgenes. H. Western blot confirming deletion of *AR*. p values for relevant differences calculated by linear regression are given in Supplementary Table 1 (induction of death by AP-2β was highly significant in all cases and there was no interaction between *TFAP2B* and the other transgenes or *AR* deletion).

**S5. Silencing of *TFAP2B* with siRNA in MDA-MB-453 cells inhibits growth and induces apoptosis.** A. Western blots showing reduced AP-2β expression with two different siRNAs at 10 nM in MDA-MB-453. B. Manual growth curves showing growth inhibition by the siRNAs used in panel A.

**S6. Peak overlap in the AP-2**β **ChIP-seq dataset.** Venn diagram showing the overlap between MACS2 peaks in the three replicates for the AP-2β ChIP-seq datasets.

**S7. AR interacts with TFAP-2β in molecular apocrine breast cancers.** A. Peptide coverage of AR and AP-2β from AR RIME experiments in MDA-MB-453 cells. Orange and pink bars represent regions in which the peptides were identified; the combined sequence coverage of identified peptides is given as a percentage. B. AP-2β unique peptides identified by RIME were tested by NCBI–Blast and five unique sequences with 100% query coverage and identity were identified (upper panel). The location of the identified peptides within the AP-2β sequence is shown in the lower panel. B. Co-immunoprecipitation (IP) of AR by antibody against AP-2β in MDA-MB-453 cells. D. Proximity ligation assay showing co-localization of AR and AP-2β in MDA-MB-453 cells grown in stripped serum then treated with either vehicle (Veh) or 10 nM DHT (left panels). E. PLA for AR and AP-2β in primary molecular apocrine breast tumours (MABC). Scale bars = 50 µm.

**S8. AP-2β, AR, FOXA1 ChIP, H3K27ac ChIP and H3K27ac HiChIP.** A-C, Apocrine genes (*CLCA2*, *KYNU*, *ERBB2*). D-F. Transcription factors (AP-2β, FOXA1, AR and GATA3). HiChIP loops are not shown for nearby peaks within the gene or overlapping the TSS.

**S9. AR, FOXA1, GATA3 and AP-2**β **motifs found in cognate peaks.** A. Sequence logos for the AR, FOXA1, GATA3 and AP-2β motifs. B. Occurrence of the motifs in A in peaks derived from ChIP for the indicated transcription factors. TP%, Estimated percentage of true positive matches. Only motifs considered to be enriched are shown, such that empty cells can be confidently assumed not to be enriched for the motif.

**S10. Overlap of published cistromes with AR+FOXA1+GATA3+AP-2**β **peaks in this study.** A&B. Overlap with peaks at active (A) and inactive (B) sites in this study. C&D. Overlap with peaks at promoters (C) and enhancers (D) in this study.

**Supplementary Table 1. Antibodies, plasmids, oligonucleotides, NGS replicates, and regression statistics for Incucyte data.** Information is grouped by sheet within the file.

