## Supplementary figures for "The transcription factor AP-2β defines active enhancers conferring molecular apocrine cell identity in breast cancer"

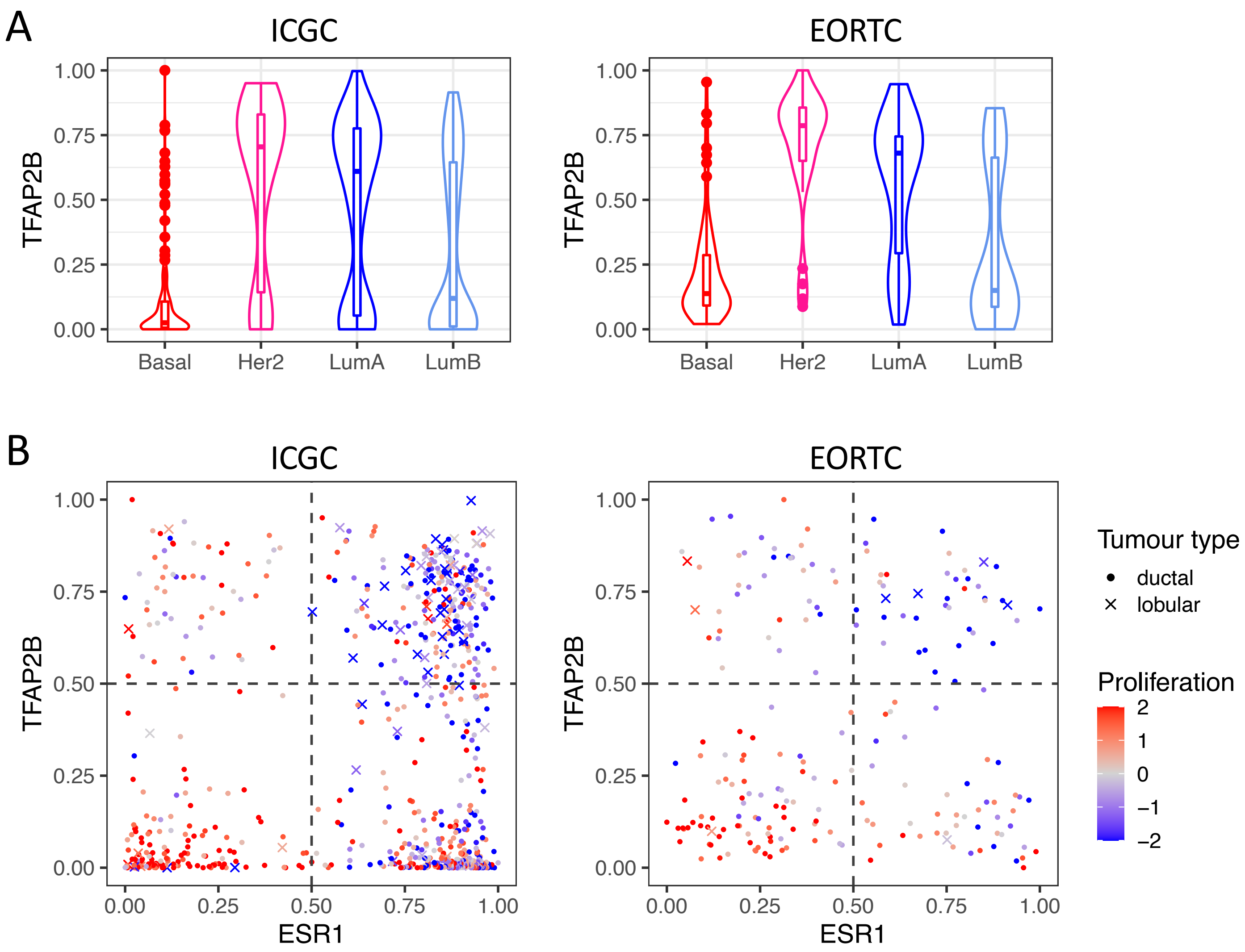

A
ICGC
EORTC
TFAP2B
1.00
0.75
0.50
0.25
0.00
Basal
Her2
LumA
LumB
B
ESR1
Tumour type
ductal
lobular
Proliferation
2
1
0
−1
−2


SF1AB

C

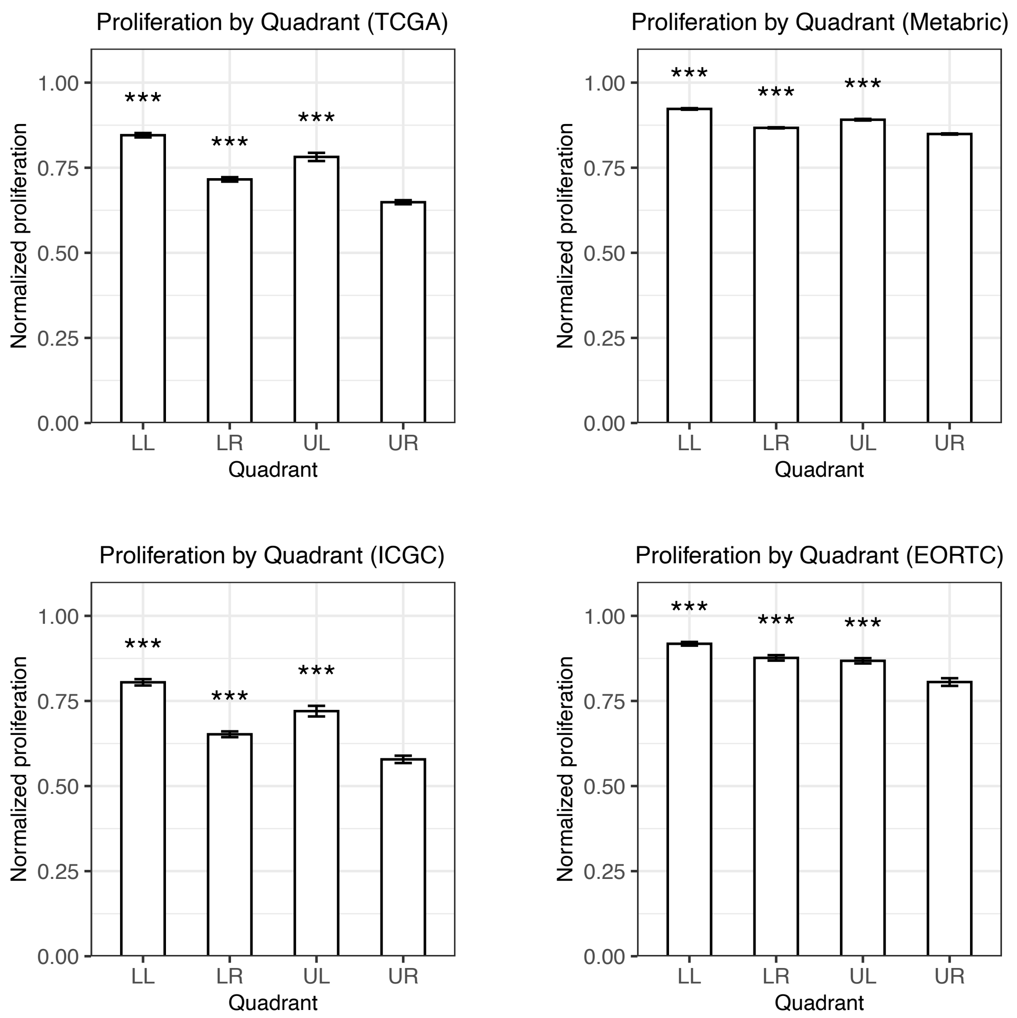

Proliferation by Quadrant (TCGA)
Proliferation by Quadrant (Metabric)
Proliferation by Quadrant (ICGC)
Proliferation by Quadrant (EORTC)
Normalized proliferation
Quadrant
LL
LR
UL
UR
***
0.00
0.25
0.50
0.75
1.00


D

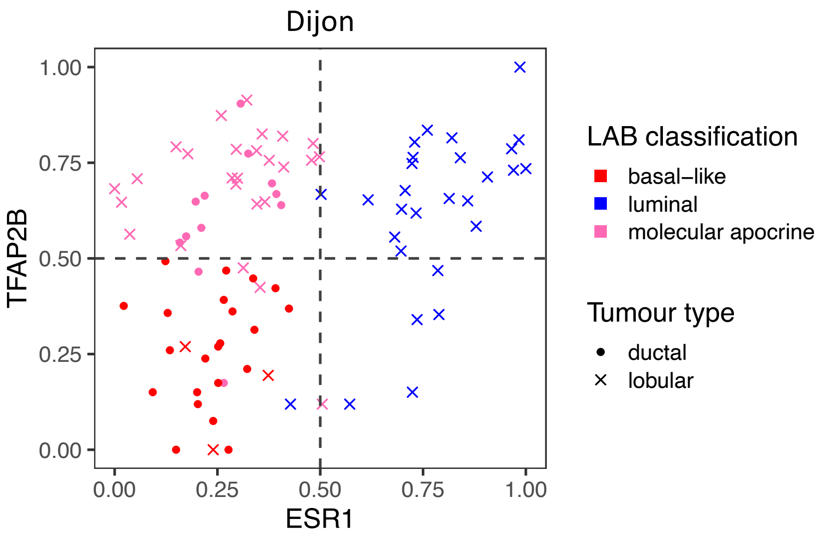

Dijon
TFAP2B
ESR1
0.00
0.25
0.50
0.75
1.00
LAB classification
basal–like
luminal
molecular apocrine
Tumour type
ductal
lobular


SF1CD

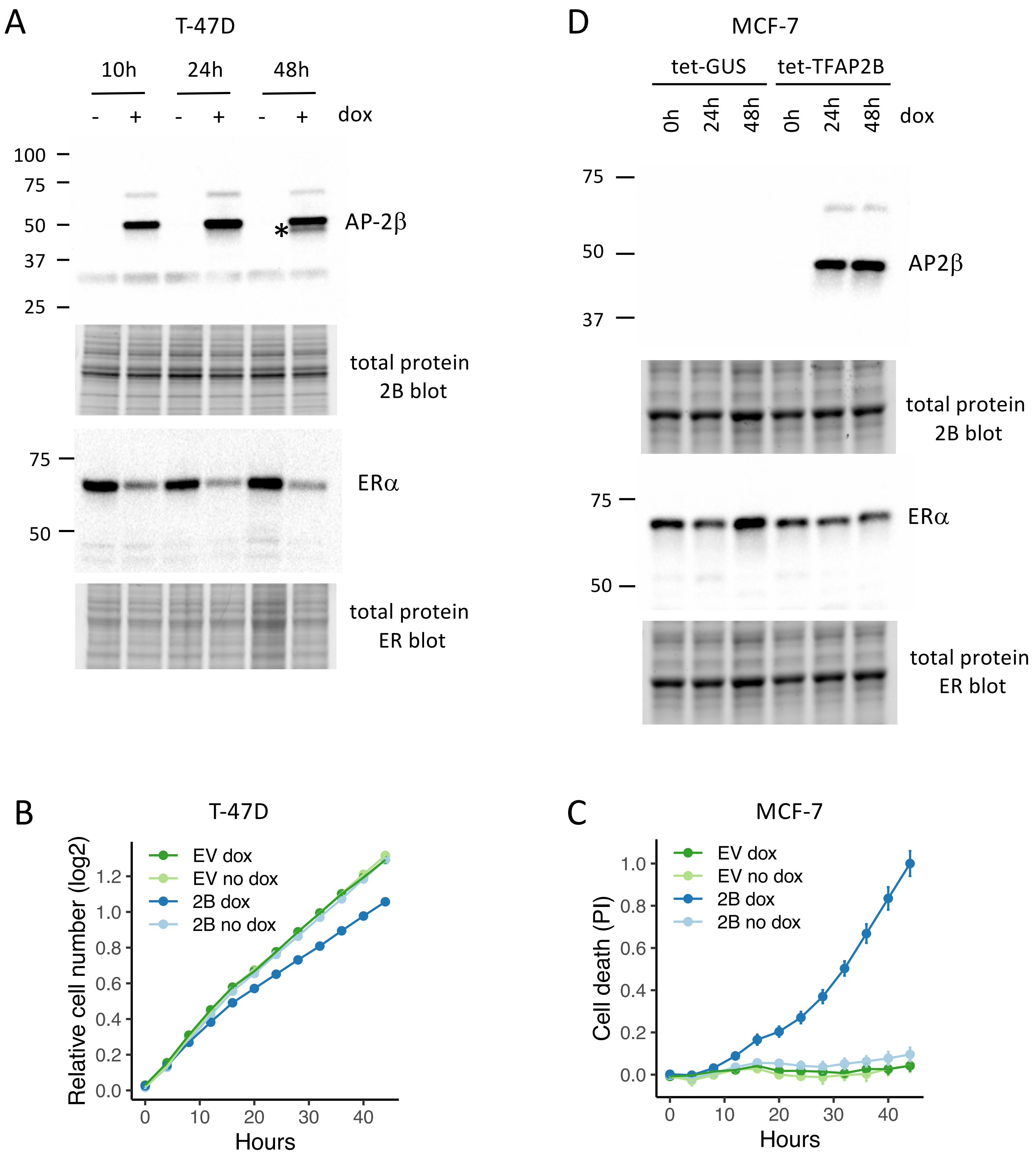
A
T-47D
10h
24h
48h
- + - + - + dox
100
75
50
37
25
AP-2β
*
total protein
2B blot
75
50
ERα
total protein
ER blot
D
MCF-7
tet-GUS
tet-TFAP2B
0h 24h 48h 0h 24h 48h dox
75
50
37
AP2β
total protein
2B blot
75
50
ERα
total protein
ER blot
B
T-47D
Relative cell number (log2)
EV dox
EV no dox
2B dox
2B no dox
1.2
1.0
0.8
0.6
0.4
0.2
0.0
0 10 20 30 40
Hours
C
MCF-7
Cell death (PI)
EV dox
EV no dox
2B dox
2B no dox
1.0
0.8
0.6
0.4
0.2
0.0
0 10 20 30 40
Hours

SF2

A

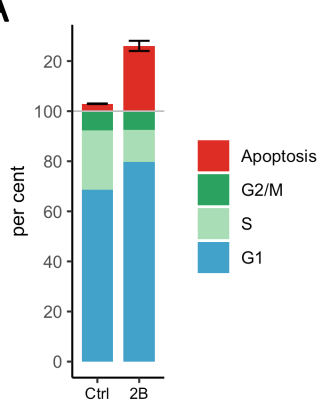


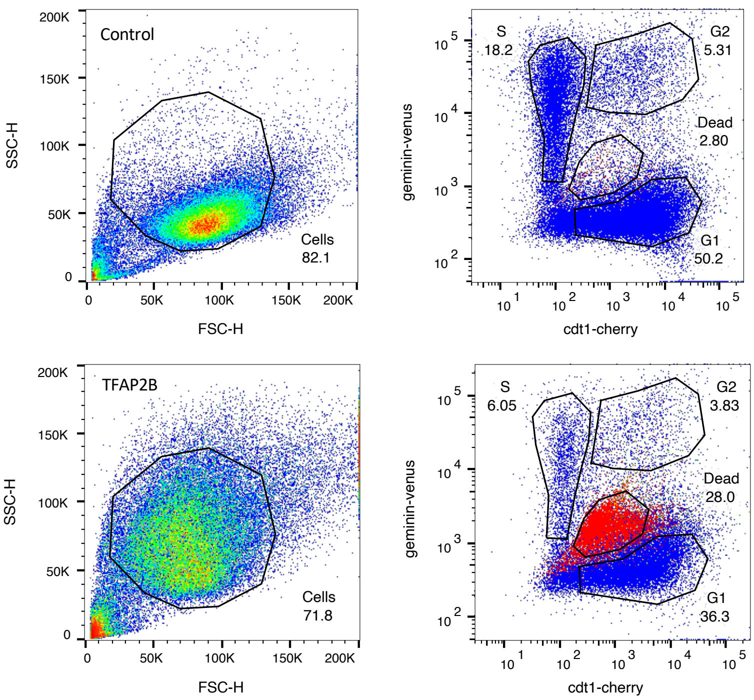


| Phase | Bonferroni p |
|---|---|
| Apoptosis | 0.0300 |
| G1 | 0.0015 |
| S | 0.0028 |
| G2/M | 1 |

B

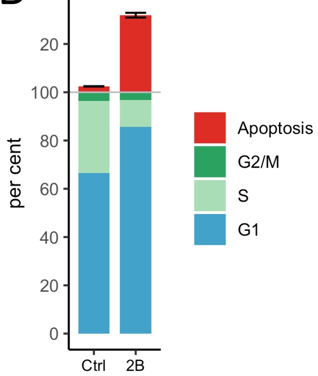


| Phase | Bonferroni p |
|---|---|
| Apoptosis | 0.0042 |
| G1 | 0.0172 |
| S | 0.0075 |
| G2/M | 1 |

C

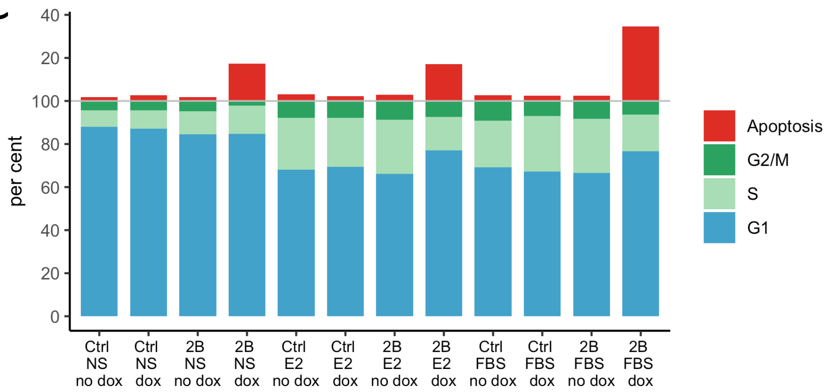


SF3

A

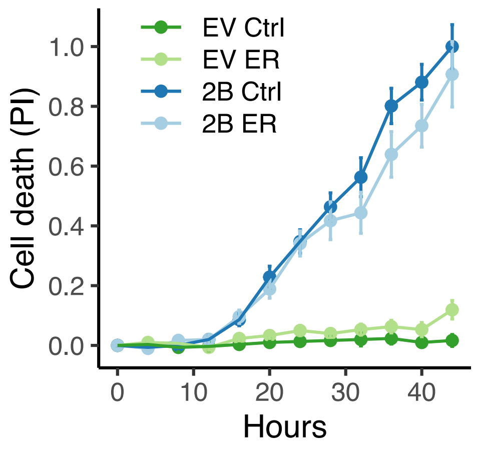
EV Ctrl
EV ER
2B Ctrl
2B ER
Cell death (PI)
1.0
0.8
0.6
0.4
0.2
0.0
0
10
20
30
40
Hours

B

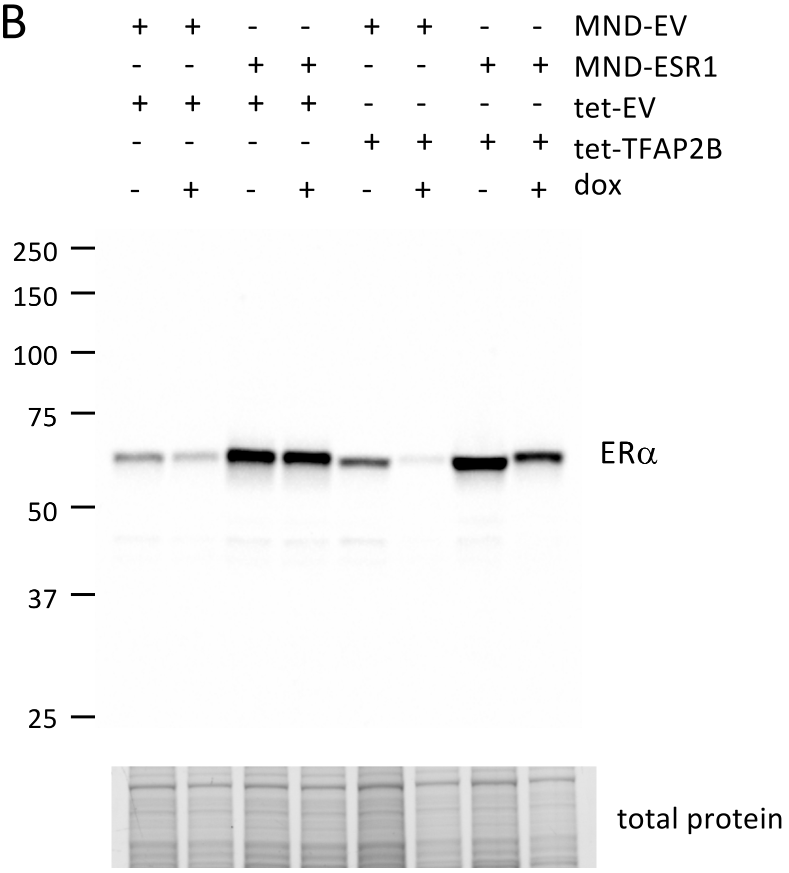
+ + - - + + - - MND-EV
- - + + - - + + MND-ESR1
+ + + + - - - - tet-EV
- - - - + + + + tet-TFAP2B
- + - + - + - + dox
250
150
100
75
50
37
25
ERα
total protein

SF4AB

C

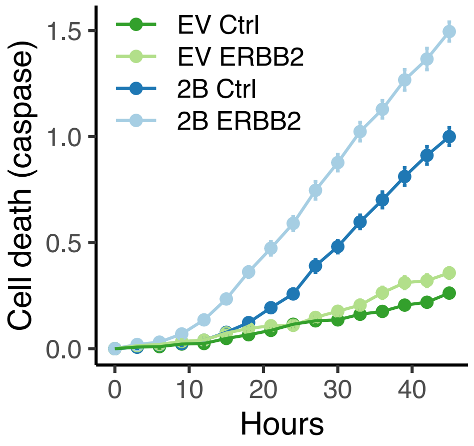
EV Ctrl
EV ERBB2
2B Ctrl
2B ERBB2
Cell death (caspase)
1.5
1.0
0.5
0.0
0
10
20
30
40
Hours

D

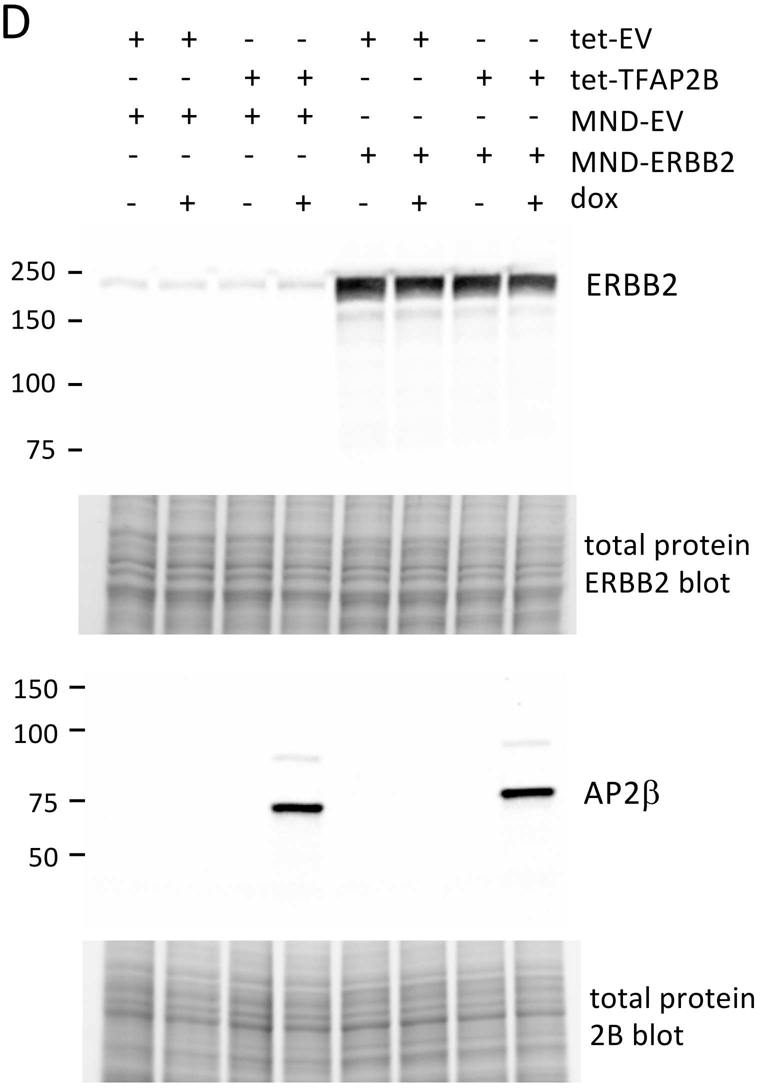
+ + - - + + - - tet-EV
- - + + - - + + tet-TFAP2B
+ + + + - - - - MND-EV
- - - - + + + + MND-ERBB2
- + - + - + - + dox
250
150
100
75
ERBB2
total protein
ERBB2 blot
150
100
75
50
AP2β
total protein
2B blot

SF4CD

E

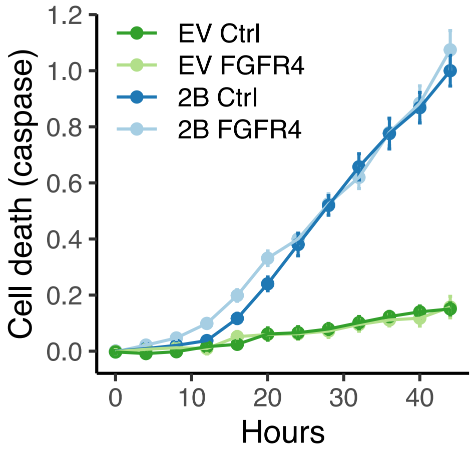

EV Ctrl
EV FGFR4
2B Ctrl
2B FGFR4
Cell death (caspase)
1.2
1.0
0.8
0.6
0.4
0.2
0.0
0
10
20
30
40
Hours


F

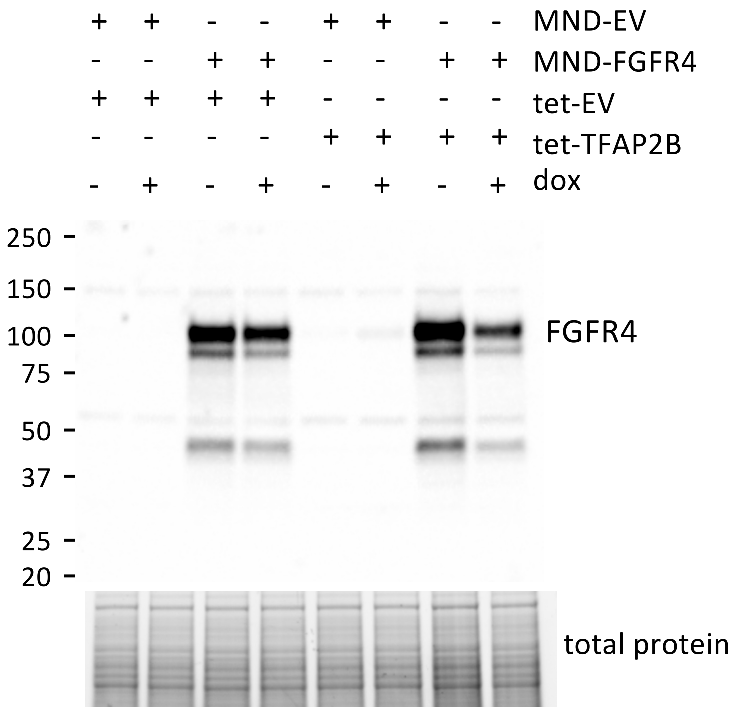

+ + - - + + - - MND-EV
- - + + - - + + MND-FGFR4
+ + + + - - - - tet-EV
- - - - + + + + tet-TFAP2B
- + - + - + - + dox
250
150
100
75
50
37
25
20
FGFR4
total protein


SF4EF

G

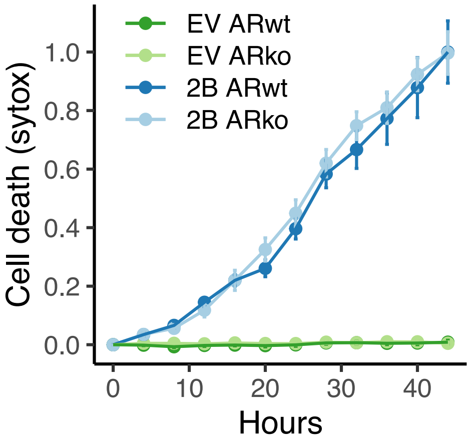
EV ARwt
EV ARko
2B ARwt
2B ARko
Cell death (sytox)
1.0
0.8
0.6
0.4
0.2
0.0
0
10
20
30
40
Hours

H

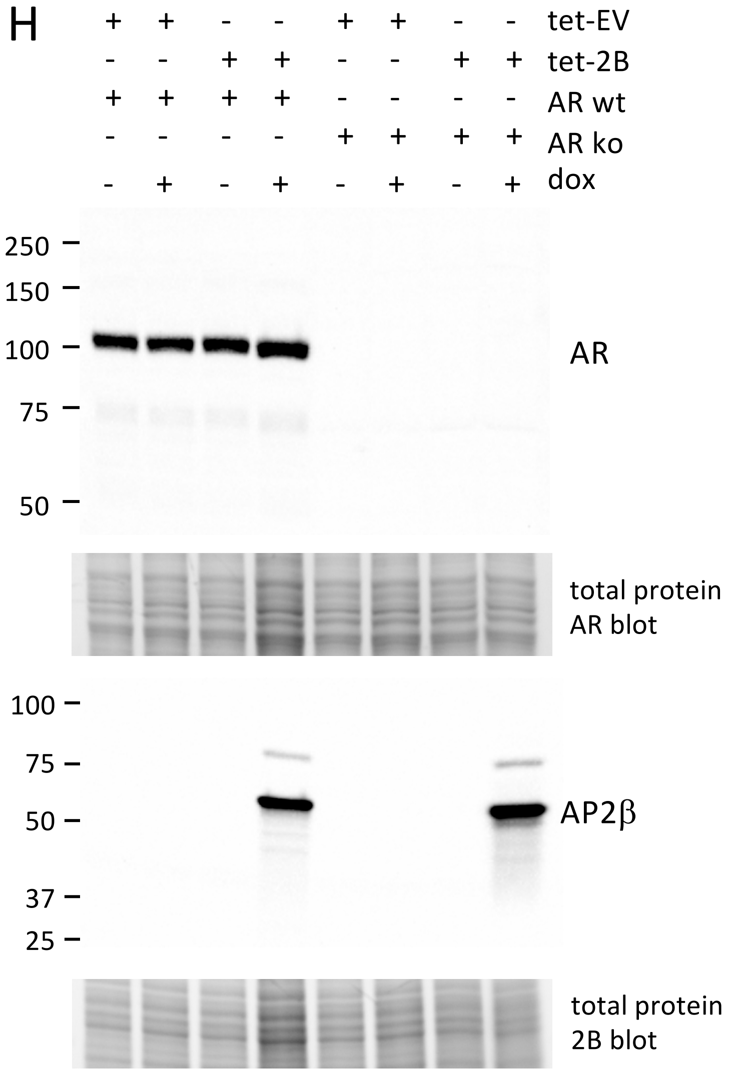
+ + - - + + - - tet-EV
- - + + - - + + tet-2B
+ + + + - - - - AR wt
- - - - + + + + AR ko
- + - + - + - + dox
250
150
100
75
50
AR
total protein
AR blot
100
75
50
37
25
AP2β
total protein
2B blot

SF4GH

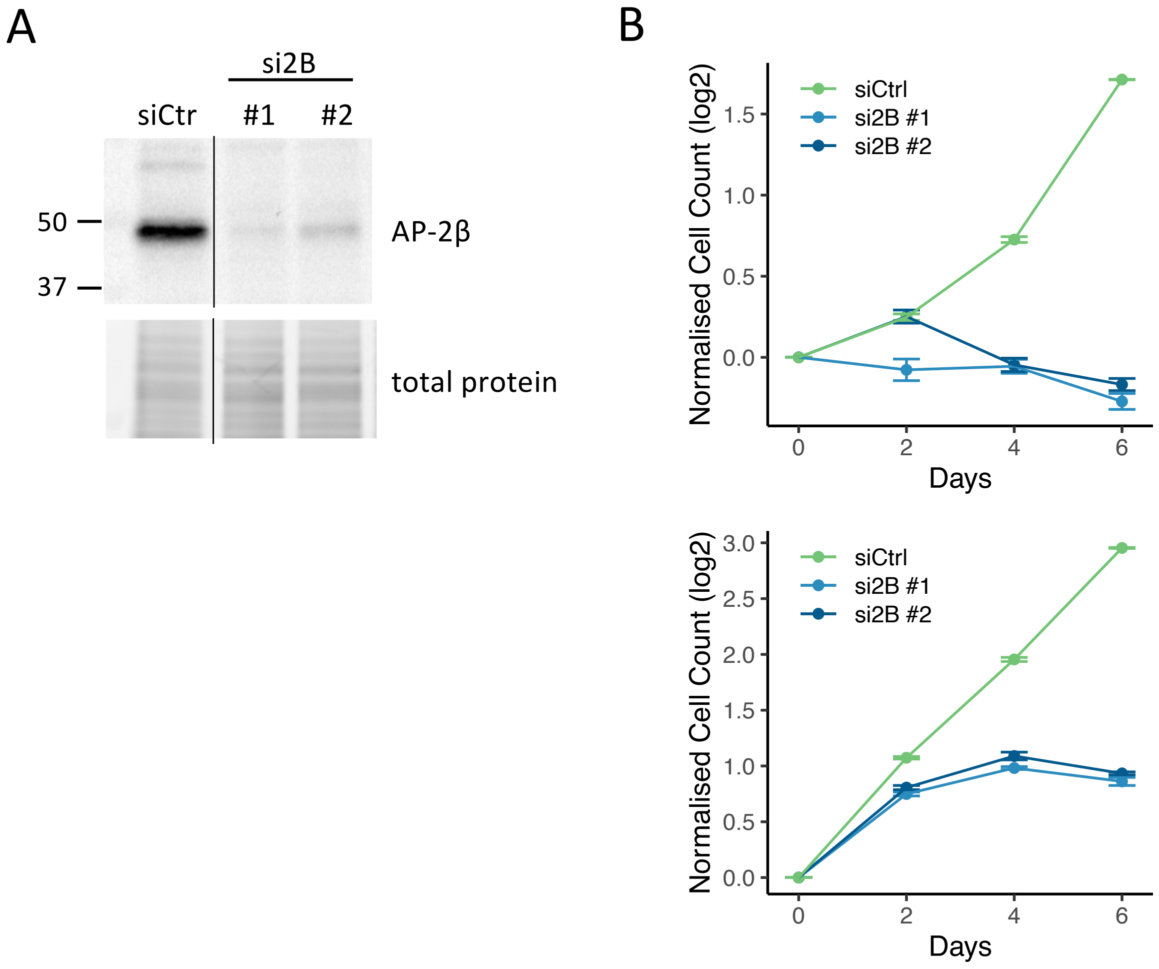


$p < 10^{-7}$ for all si2B vs siCtrl differences on day 6 (ANOVA + Dunnett)

SF5

TFAP2B ChIP-seq peaks

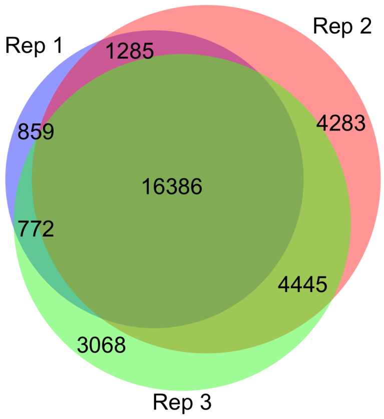


SF6

A

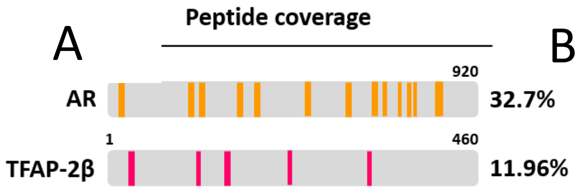


B

| TFAP-2β unique peptides | Query cover | Identities (%) | E value |
|---|---|---|---|
| SVTSLMMNK | 100% | 100% | 3.00E-04 |
| AVSEYLNR | 100% | 100% | 0.008 |
| AALPQLSGLDPR | 100% | 100% | 3.00E-06 |
| LVENVKYEDIYEDRHDGVPSHSSR | 100% | 100% | 3.00E-20 |
| QRQEVGSEAGSLLPQPR | 100% | 100% | 2.00E-11 |

```
  1 MHSPPRDQAA IMLWKLVENV KYEDIYEDRH DGVPSHSSRL SQLGSVSQGP YSSAPPLSHT
 61 PSSDFQPPYF PPPYQPLPYH QSQDPYSHVN DPYSLNPLHQ PQQHPWGQRQ RQEVGSEAGS
121 LLPQPRAALP QLSGLDPRRD YHSVRRPDVL LHSAHHGLDA GMGDSLSLHG LGHPGMEDVQ
181 SVEDANNSGM NLLDQSVIKK VPVPPKSVTS LMMNKDGFLG GMSVNTGEVF CSVPGRLSLL
241 SSTSKYKVTV GEVQRRLSPP ECLNASLLGG VLRRAKSKNG GRSLRERLEK IGLNLPAGRR
301 KAANVTLLTS LVEGEAVHLA RDFGYICETE FPAKAVSEYL NRQHTDPSDL HSRKNMLLAT
361 KQLCKEFTDL LAQDRTPIGN SRPSPILEPG IQSCLTHFSL ITHGFGAPAI CAALTALQNY
421 LTEALKGMDK MFLNNTTTNR HTSGEGPGSK TGDKEEKHRK
```

C

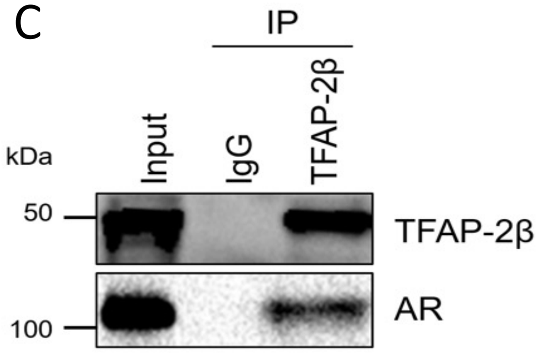


D

E

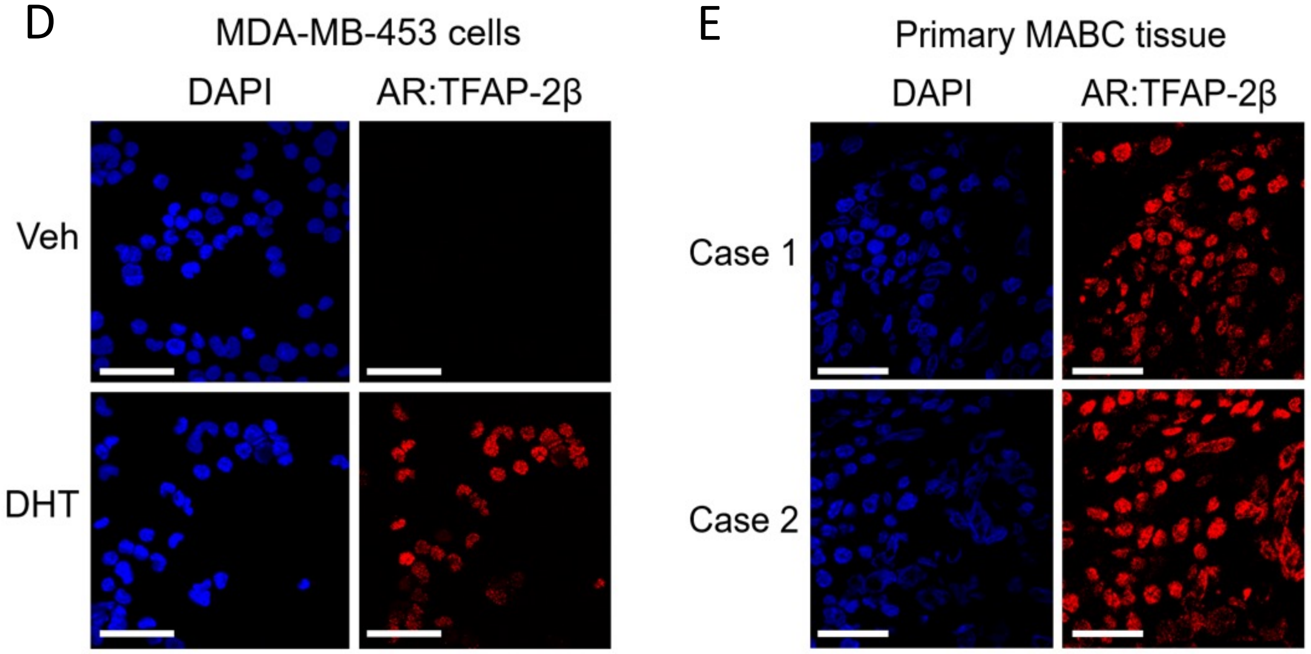

A

B

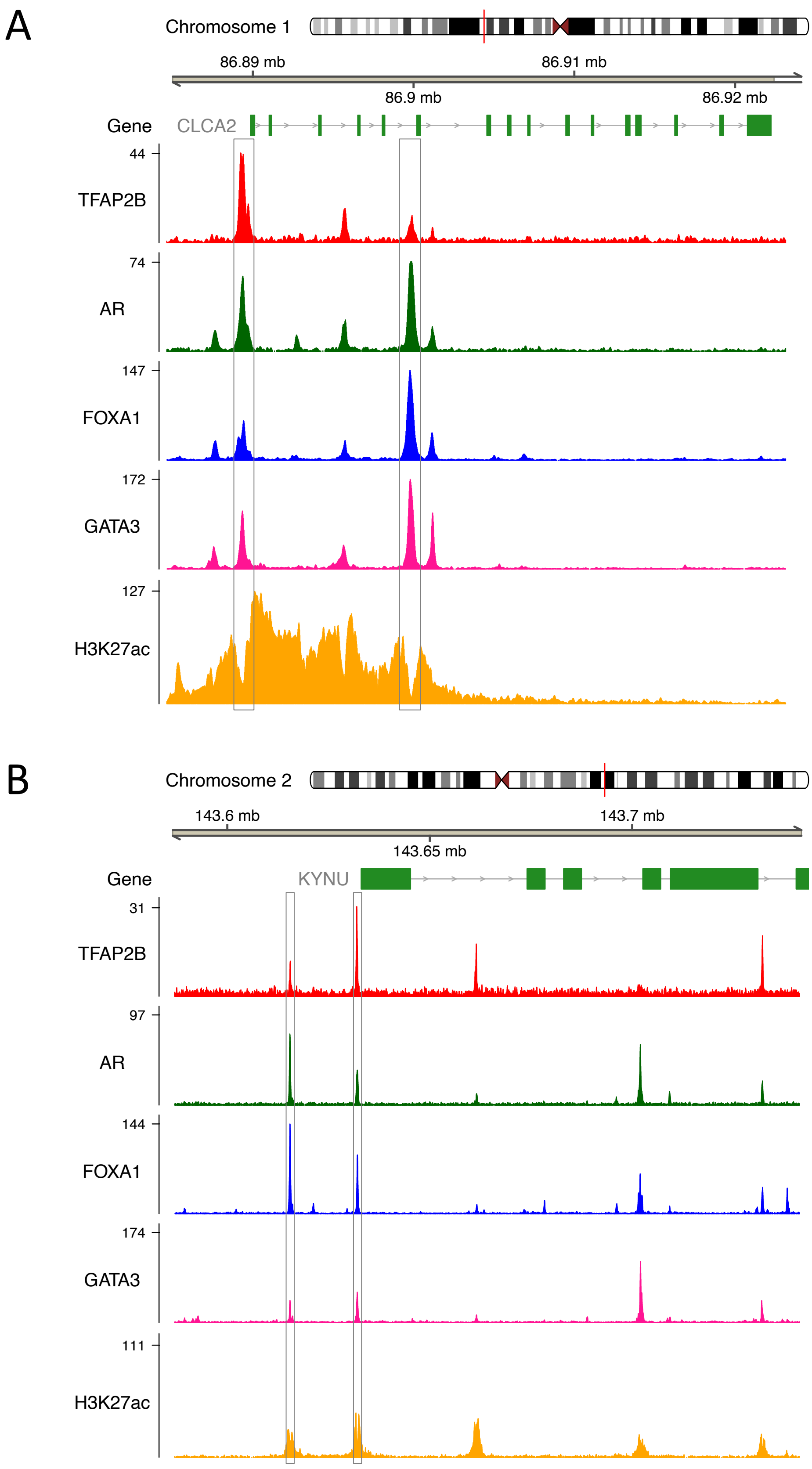

Chromosome 1
86.89 mb
86.91 mb
86.9 mb
86.92 mb
Gene
CLCA2
44
TFAP2B
74
AR
147
FOXA1
172
GATA3
127
H3K27ac
Chromosome 2
143.6 mb
143.7 mb
143.65 mb
Gene
KYNU
31
TFAP2B
97
AR
144
FOXA1
174
GATA3
111
H3K27ac


SF8AB

C

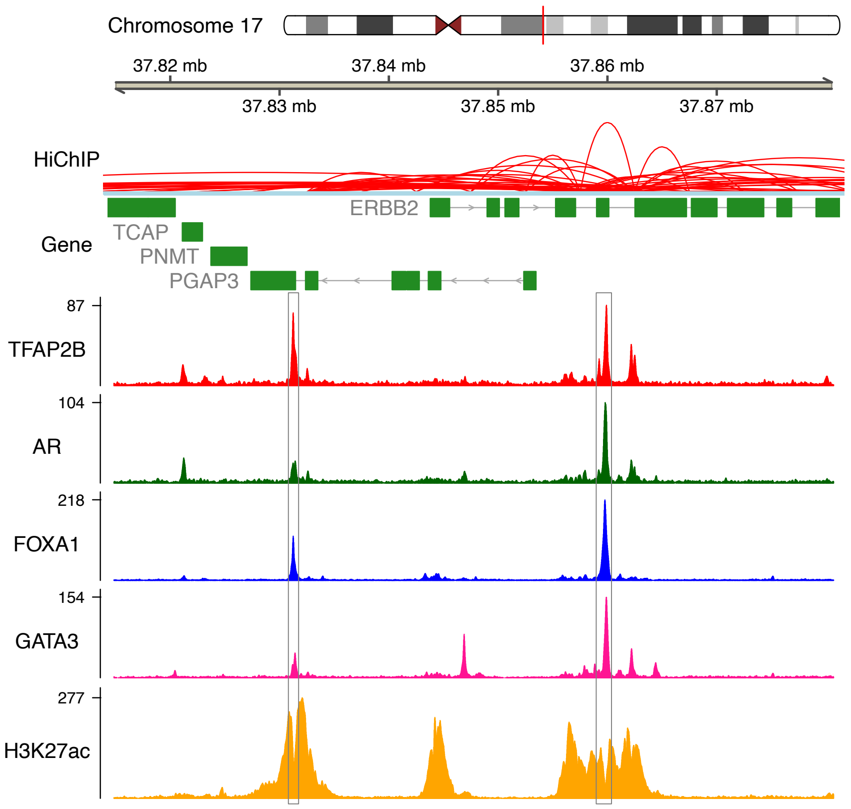

Chromosome 17
37.82 mb
37.84 mb
37.86 mb
37.83 mb
37.85 mb
37.87 mb
HiChIP
ERBB2
TCAP
Gene
PNMT
PGAP3
87
TFAP2B
104
AR
218
FOXA1
154
GATA3
277
H3K27ac


D

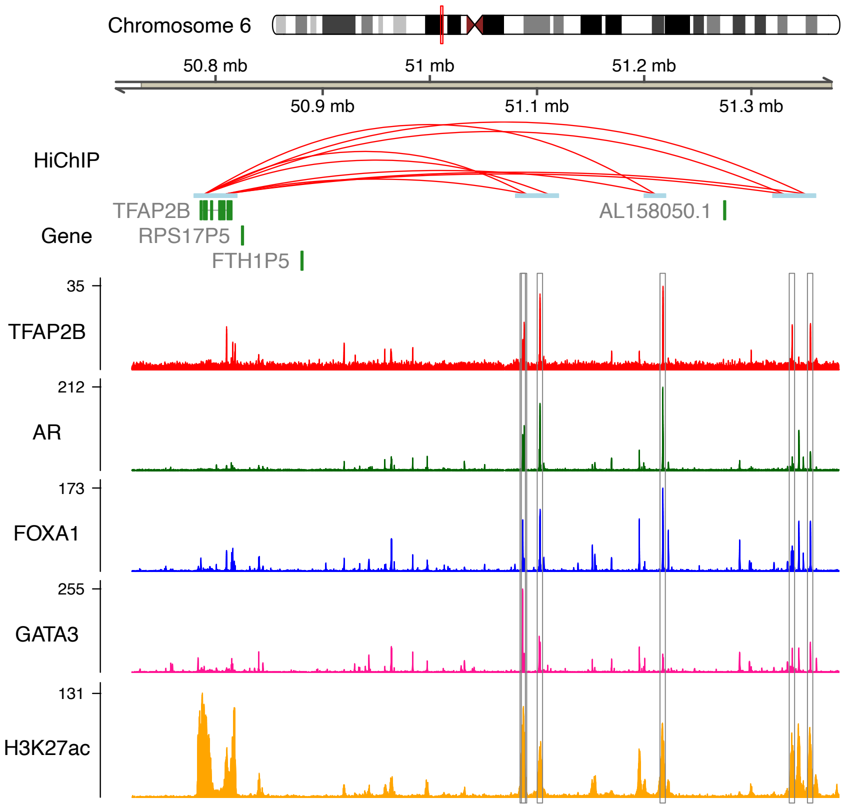

Chromosome 6
50.8 mb
51 mb
51.2 mb
50.9 mb
51.1 mb
51.3 mb
HiChIP
TFAP2B
AL158050.1
Gene
RPS17P5
FTH1P5
35
TFAP2B
212
AR
173
FOXA1
255
GATA3
131
H3K27ac


SF8CD

E

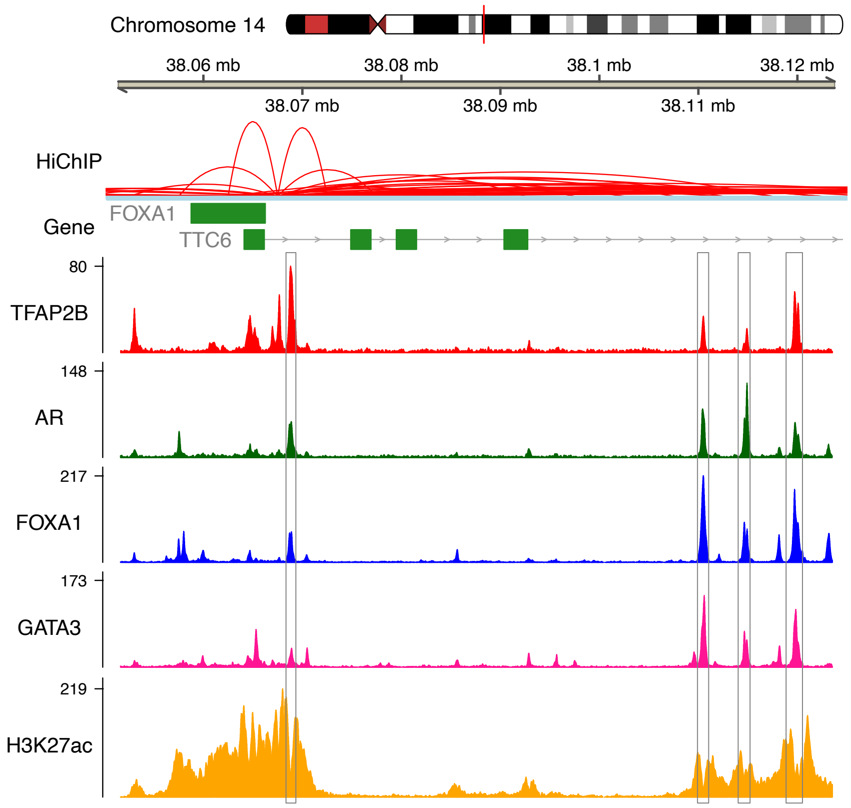

Chromosome 14
38.06 mb
38.08 mb
38.1 mb
38.12 mb
38.07 mb
38.09 mb
38.11 mb
HiChIP
Gene
FOXA1
TTC6
80
TFAP2B
148
AR
217
FOXA1
173
GATA3
219
H3K27ac


F

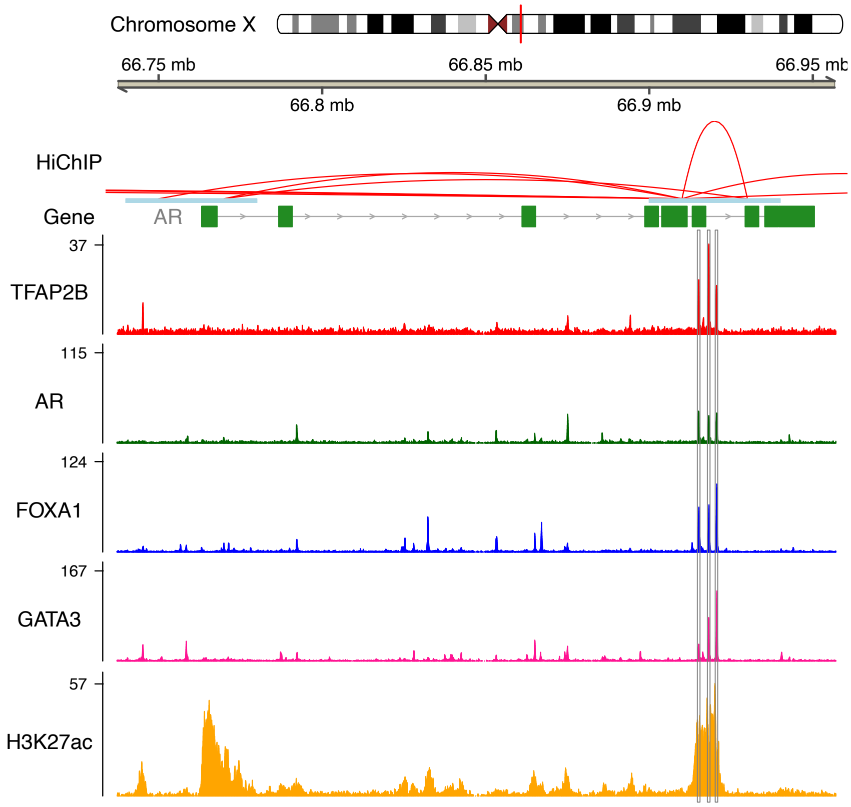

Chromosome X
66.75 mb
66.85 mb
66.95 mb
66.8 mb
66.9 mb
HiChIP
Gene
AR
37
TFAP2B
115
AR
124
FOXA1
167
GATA3
57
H3K27ac


SF8EF

G

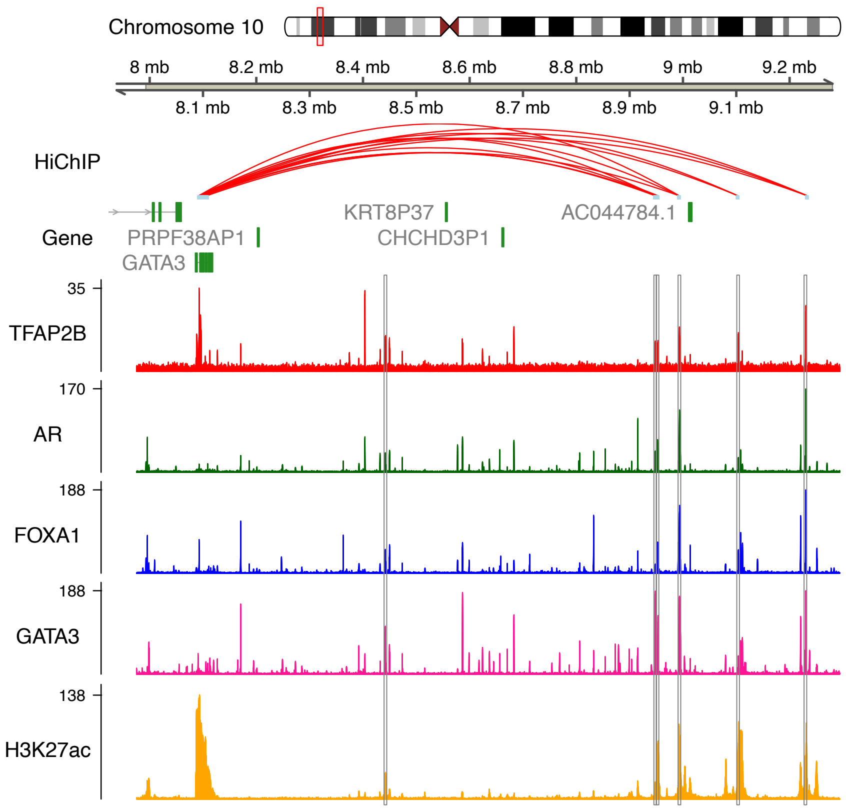

Chromosome 10
8 mb
8.2 mb
8.4 mb
8.6 mb
8.8 mb
9 mb
9.2 mb
8.1 mb
8.3 mb
8.5 mb
8.7 mb
8.9 mb
9.1 mb
HiChIP
Gene
KRT8P37
AC044784.1
PRPF38AP1
CHCHD3P1
GATA3
35
TFAP2B
170
AR
188
FOXA1
188
GATA3
138
H3K27ac


SF8G

A

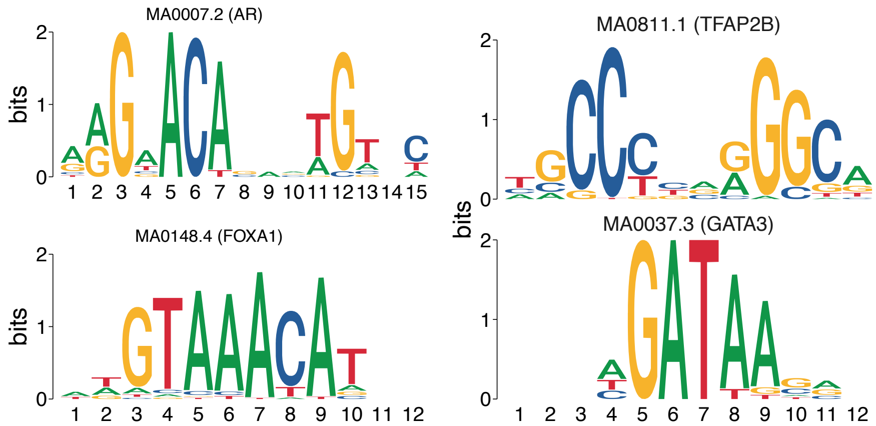
MA0007.2 (AR)
MA0811.1 (TFAP2B)
MA0148.4 (FOXA1)
MA0037.3 (GATA3)
bits
0
1
2

B

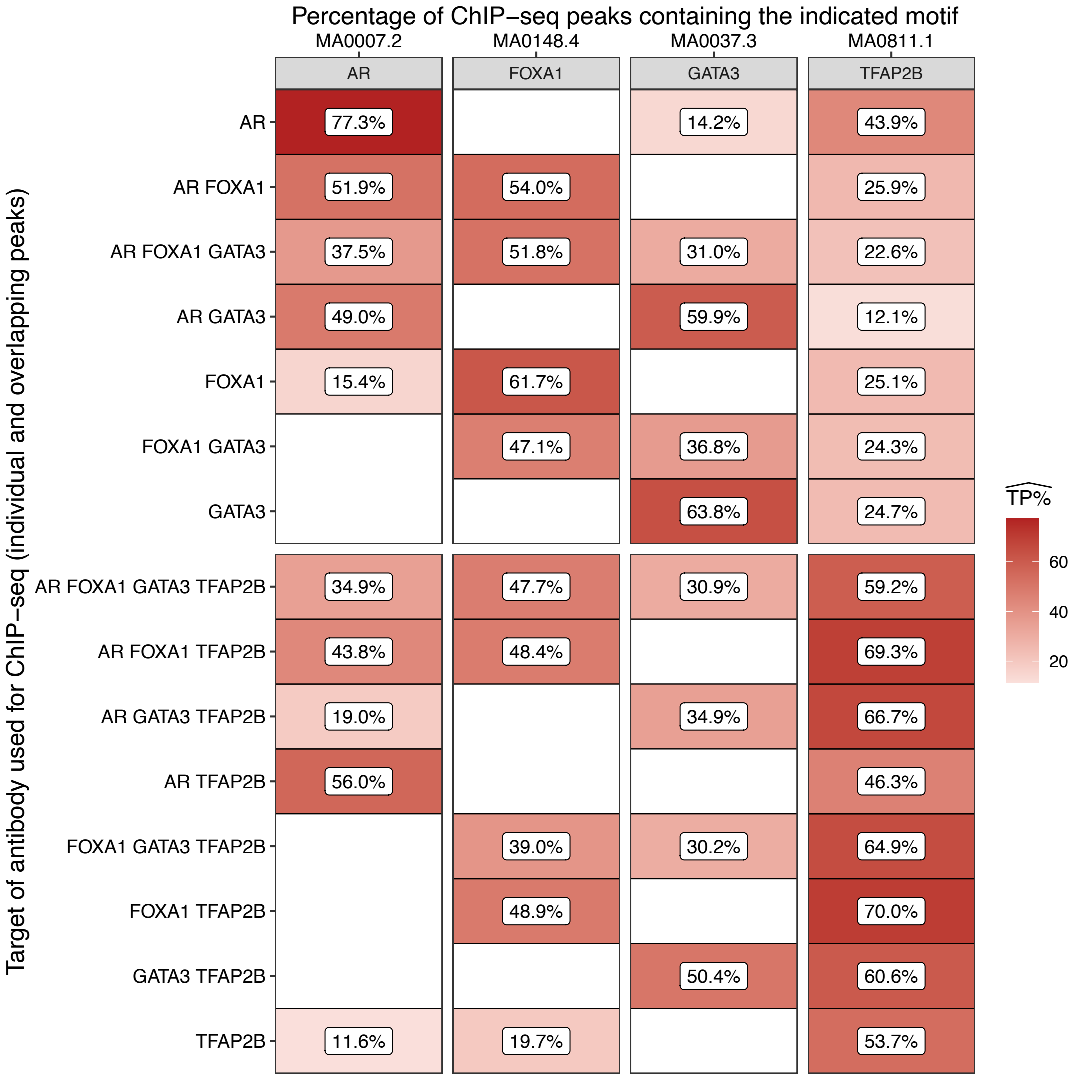
Percentage of ChIP−seq peaks containing the indicated motif
MA0007.2
MA0148.4
MA0037.3
MA0811.1
AR
FOXA1
GATA3
TFAP2B
Target of antibody used for ChIP−seq (individual and overlapping peaks)
AR 77.3% 14.2% 43.9%
AR FOXA1 51.9% 54.0% 25.9%
AR FOXA1 GATA3 37.5% 51.8% 31.0% 22.6%
AR GATA3 49.0% 59.9% 12.1%
FOXA1 15.4% 61.7% 25.1%
FOXA1 GATA3 47.1% 36.8% 24.3%
GATA3 63.8% 24.7%
AR FOXA1 GATA3 TFAP2B 34.9% 47.7% 30.9% 59.2%
AR FOXA1 TFAP2B 43.8% 48.4% 69.3%
AR GATA3 TFAP2B 19.0% 34.9% 66.7%
AR TFAP2B 56.0% 46.3%
FOXA1 GATA3 TFAP2B 39.0% 30.2% 64.9%
FOXA1 TFAP2B 48.9% 70.0%
GATA3 TFAP2B 50.4% 60.6%
TFAP2B 11.6% 19.7% 53.7%
TP%
60
40
20

SF9

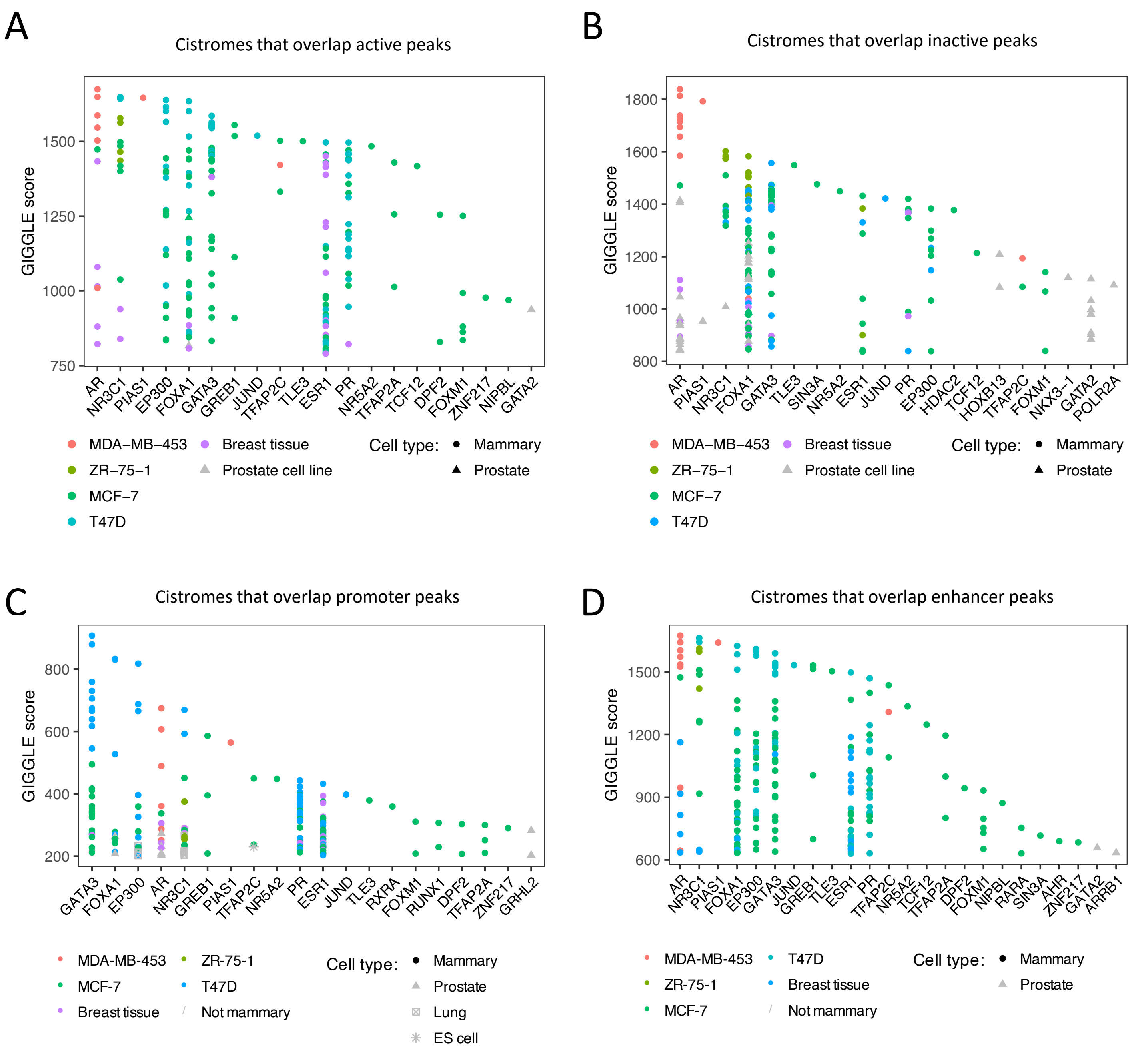
A
Cistromes that overlap active peaks
GIGGLE score
750
1000
1250
1500
AR
NR3C1
PIAS1
EP300
FOXA1
GATA3
GREB1
JUND
TFAP2C
TLE3
ESR1
PR
NR5A2
TFAP2A
TCF12
DPF2
FOXM1
ZNF217
NIPBL
GATA2
MDA−MB−453
ZR−75−1
MCF−7
T47D
Breast tissue
Prostate cell line
Cell type:
Mammary
Prostate
B
Cistromes that overlap inactive peaks
GIGGLE score
800
1000
1200
1400
1600
1800
AR
PIAS1
NR3C1
FOXA1
GATA3
TLE3
SIN3A
NR5A2
ESR1
JUND
PR
EP300
HDAC2
TCF12
HOXB13
TFAP2C
FOXM1
NKX3−1
GATA2
POLR2A
MDA−MB−453
ZR−75−1
MCF−7
T47D
Breast tissue
Prostate cell line
Cell type:
Mammary
Prostate
C
Cistromes that overlap promoter peaks
GIGGLE score
200
400
600
800
GATA3
FOXA1
EP300
AR
NR3C1
GREB1
PIAS1
TFAP2C
NR5A2
PR
ESR1
JUND
TLE3
RXRA
FOXM1
RUNX1
DPF2
TFAP2A
ZNF217
GRHL2
MDA-MB-453
MCF-7
Breast tissue
ZR-75-1
T47D
Not mammary
Cell type:
Mammary
Prostate
Lung
ES cell
D
Cistromes that overlap enhancer peaks
GIGGLE score
600
900
1200
1500
AR
NR3C1
PIAS1
FOXA1
EP300
GATA3
JUND
GREB1
TLE3
ESR1
PR
TFAP2C
NR5A2
TCF12
TFAP2A
DPF2
FOXM1
NIPBL
RARA
SIN3A
AHR
ZNF217
GATA2
ARRB1
MDA-MB-453
ZR-75-1
MCF-7
T47D
Breast tissue
Not mammary
Cell type:
Mammary
Prostate

SF10
